# Damaging mutations in LXRα uncouple lipogenesis from hepatotoxicity and implicate hepatic cholesterol sensing in human liver health

**DOI:** 10.1101/2024.04.28.591512

**Authors:** Sam M Lockhart, Milan Muso, Ilona Zvetkova, Brian YH Lam, Alessandra Ferrari, Erik Schoenmakers, Katie Duckett, Jack Leslie, Beatriz Romartínez-Alonso, John Tadross, Raina Jia, Eugene J. Gardner, Katherine Kentistou, Yajie Zhao, Felix Day, Alexander Mörseburg, Kara Rainbow, Debra Rimmington, Matteo Mastantuoni, James Harrison, Meritxell Nus, Khalid Guma’a, Sam Sherratt-Mayhew, Xiao Jiang, Katherine R Smith, Dirk S Paul, Ben Jenkins, Albert Koulman, Maik Pietzner, Claudia Langenberg, Nick Wareham, Giles S Yeo, Krishna Chatterjee, John Schwabe, Fiona Oakley, Derek Mann, Peter Tontonoz, Tony Coll, Ken Ong, John R.B. Perry, Stephen O’Rahilly

**Affiliations:** Medical Research Council (MRC) Metabolic Diseases Unit, Institute of Metabolic Science, University of Cambridge, Cambridge, UK; Department of Pathology and Laboratory Medicine, University of California, Los Angeles; CA 90095, USA; Newcastle University Centre for Cancer, Newcastle University, Newcastle upon Tyne, UK; Newcastle Fibrosis Research Group, Bioscience Institute, Faculty of Medical Sciences, Newcastle University, Newcastle upon Tyne, UK; Institute of Structural and Chemical Biology, Department of Molecular and Cell Biology, University of Leicester, Leicester, United Kingdom; Medical Research Council (MRC) Epidemiology Unit, Institute of Metabolic Science, University of Cambridge, Cambridge, UK; VPD Heart and Lung Research Institute, Dept. Medicine, University of Cambridge, Cambridge, UK; Centre for Genomics Research, Discovery Sciences, BioPharmaceuticals R&D, AstraZeneca, Cambridge, UK; NIHR BRC Core Metabolomics and Lipidomics Laboratory, Metabolic Research Laboratories, Institute of Metabolic Science, University of Cambridge, Cambridge CB2 1GG, UK; Computational Medicine, Berlin Institute of Health at Charité - Universitätsmedizin Berlin, Berlin, Germany; Precision Healthcare University Research Institute, Queen Mary University of London, London, UK; NIHR Cambridge Biomedical Research Centre, Cambridge, UK

## Abstract

The nuclear receptor Liver X Receptor-α (LXRα) activates lipogenic gene expression in hepatocytes. Its inhibition has therefore been proposed as a strategy to treat metabolic-dysfunction-associated steatotic liver disease (MASLD). In order to understand the impact of reducing LXRα activity on human health we first examined the association between the carriage of rare loss of function mutations in *NR1H3* (encoding LXRα) and metabolic and hepatic phenotypes. We identified 63 rare predicted damaging variants in the ligand binding domain of LXRα in 454,787 participants in UK Biobank. On functional characterisation, 42 of these were found to be severely impaired. Consistent with loss of the lipogenic actions of LXRα, carriers of damaging mutations in LXRα had reduced serum triglycerides (ß=-0.13 s.d. ±0.03, P=2.7x10^-5^, N(carriers)=971). Surprisingly, these carriers also had elevated concentrations of serum liver enzymes (e.g. ALT: ß=0.17s.d. ±0.03, P=1.1x10^-8^, N(carriers)=972) with a 35% increased risk of clinically significant elevations in ALT (OR=1.32, 95%CI:1.15-1.53, P=1.2x10^-4^, N(carriers)=972), suggestive of hepatotoxicity. We generated a knock-in mouse carrying one of the most severely damaging mutations (*Nr1h3* p.W441R) which we demonstrated to have dominant negative properties. Homozygous knock-in mice rapidly developed severe hepatitis and fibrotic liver injury following exposure to western diet despite markedly reduced steatosis, liver triglycerides and lipogenic gene expression. This phenotype was completely rescued by viral over-expression of wildtype LXRα specifically in hepatocytes, indicating a cell-autonomous effect of the mutant on hepatocyte health. While homozygous LXRα knockout mice showed some evidence of hepatocyte injury under similar dietary conditions, the phenotype of the LXRα^W441R/W441R^ mouse was much more severe, suggesting that dominant negative mutations that actively co-repress target genes can result in pathological impacts significantly more severe than those seen with simple absence of the receptor. In summary, our results show that loss of function mutations in LXRα occur in at least 1/450 people and are associated with evidence of liver dysfunction. These findings implicate LXRα in the maintenance of human liver health, identify a new murine model of rapidly progressive fibrotic liver disease and caution against LXR antagonism as a therapeutic strategy for MASLD.

## Introduction

Liver-X-Receptors (LXR) are oxysterol-regulated nuclear receptors critical for cellular and organismal cholesterol homeostasis [1]. Oxysterols, generated as a consequence of rising intracellular cholesterol levels, bind to the ligand binding domain (LBD) of LXR, resulting in a conformational change that facilitates co-repressor dissociation, co-activator recruitment, and transcriptional activation [2–5]. In hepatocytes, LXRα is the dominant isoform and its activation results in a transcriptional programme that normalises cellular cholesterol via modulation of cholesterol biosynthesis, uptake and excretion [1]. For example, biliary cholesterol excretion is enhanced via induction of the cholesterol efflux transporters *ABCG5* and *ABCG8* and upregulation of the E3 ubiquitin ligase RNF145, which targets and downregulates proteins participating in cholesterol biosynthesis [6–9]. LXR agonism also activates reverse cholesterol transport through induction of *ABCG1* and *ABCA1* in the periphery resulting in cholesterol efflux to HDL for transport to the liver [10–12]. An additional consequence of LXRα activation is upregulation of hepatic lipogenesis via induction of the lipogenic factors *SREBP1c*, *FASN* and *SCD1*, amongst others [1]. Thus, in mice, synthetic LXR agonists result in an increase in liver and serum triglycerides that appears largely dependent on hepatocyte LXRα [10, 13]. In addition, pharmacological LXR-agonism exhibits species-dependent effects on lipoprotein metabolism. LXR-agonists increase low density lipoprotein (LDL) cholesterol in higher primates, likely via induction of the ubiquitin-ligase MYLIP which reduces hepatocyte surface expression of LDLR and impairs LDL cholesterol clearance from the serum [14, 15]. LXR agonists also reduce HDL cholesterol in humans and non-human primates, possibly via increases in CETP, which mice do not naturally express [15, 16]. Studies in mice have also implicated LXRs in regulation of glucose homeostasis [17, 18].

Initial efforts to translate LXR biology to clinical application focused on using LXR-agonism to enhance reverse cholesterol transport to prevent atherosclerotic cardiovascular disease [1, 15, 19]. However, these efforts have been frustrated by challenges in dissociating the effects of agonism on reverse cholesterol transport from the undesirable effects on hepatic lipogenesis and by species-related effects on lipoprotein metabolism [14, 15]. Recently, interest has grown in adopting the converse approach: inhibiting hepatic LXR to suppress lipogenesis and hepatocyte triglyceride accumulation in the context of MASLD, a condition characterised by hepatic steatosis and cardiometabolic risk factors without significant alcohol consumption [20]. In a proportion of individuals, neutral lipid can be stored in the liver without significant hepatic inflammation. In a minority, steatosis is associated with the development of inflammation, so-called metabolic associated steatohepatitis (MASH), which can result in fibrosis, cirrhosis and end stage liver disease [21, 22]. While the mechanistic basis of this heterogeneity is not clear, abundance of triglyceride in liver likely plays a causal role; coding variants which increase liver fat via various mechanisms increase risk of liver disease and cirrhosis [23–25], and rare, predicted loss of function mutations in a lipid droplet binding protein CIDEB reduce liver fat, the risk of liver disease and cirrhosis [23].

It is therefore reasonable to ask whether suppression of lipogenesis via inverse agonism of LXR might be beneficial in MASLD. Indeed, treatment with LXR-inverse agonists reduce steatosis, inflammation and fibrosis in rodent models [20, 26, 27] and early-stage clinical trials of LXR-inverse agonists for hypertriglyceridaemia and MASLD are in progress [28]. However, increased hepatic cholesterol has been observed in MASLD and is a putative risk factor for MASH [29–31]. Moreover, several mouse models with hepatic cholesterol accumulation due to genetic perturbation of key regulators of cholesterol homeostasis (including LXRα) exhibit evidence of liver injury [32–35]. Increasing dietary cholesterol content increases the hepatic inflammation and fibrosis associated with high fat diets in mice [36, 37] and has been shown to stimulate a fibrogenic pathway in liver via TAZ-dependent activation of Indian hedgehog pathway [38]. Thus, impaired hepatic cholesterol clearance may undermine the therapeutic utility of LXR inverse agonists in MASLD.

It is increasingly recognised that human genetics can assist drug discovery by the validation of drug targets and the prediction of unwanted effects of target engagement [39, 40]. Given the translational interest in LXR in MASLD and other facets of cardiometabolic health we sought to determine the effects of damaging mutations in LXRα, the dominant LXR isoform in the liver, on human cardiometabolic health. We present a detailed assessment of the functional consequences of rare mutations in the ligand-binding domain of LXRα, identifying loss of function, gain of function and dominant negative mutations. We delineate the phenotypic consequences of these mutations in human participants of UK Biobank demonstrating a novel dose response relationship between LXR-activity and serum HDL cholesterol concentration and provide evidence of a hepatotoxic effect of LXR-haploinsubfficiency in humans despite expected reduction in serum triglycerides and triglycerides in VLDL – consistent with impaired hepatic lipogenesis. We demonstrate that knock-in mice carrying a damaging, dominant negative mutation in LXRα develop liver inflammation and severe fibrotic liver injury despite marked reduction in liver triglycerides and steatosis. Together our results highlight the essentiality of intact LXRα-signalling and hepatic cholesterol homeostasis for liver health.

## Results

### Identification of naturally occurring damaging mutations in the ligand binding domain of LXRα

To identify human carriers of damaging mutations in LXRα we interrogated exome sequencing data from 454,756 UKBB participants. We focused on the ligand binding domain as experience with naturally occurring mutations in other nuclear receptors (e.g. PPARγ, TRα/β) indicates a strong enrichment of pathological variants in this region [41–43]. We selected (see methods) a total of 63 rare variants (MAF<0.1%, 57 predicted deleterious missense variants and 6 protein truncating variants (PTVs) near the C-Terminus, all present in heterozygosity) in the ligand binding domain of LXRα (Supplementary Table 1) for characterisation in two different assays of nuclear receptor function: i) A transactivation assay using an LXR-luciferase reporter to monitor LXR-activity in response to increasing concentrations of the synthetic LXR-agonist T0901317, ii) a co-expression assay examining the ability of candidate LXRα variants to impair co-transfected wildtype LXRα as an index of dominant negative activity. We identified 23 mutations with evidence of dominant negativity (DN: N(carriers)=162, cumulative MAF=0.04%), 20 mutations with evidence of impaired transactivation without significant dominant negativity (LOF: N(carriers)=642, cumulative MAF=0.14%) and 4 gain of function mutations (GOF: N(carriers)=52, cumulative MAF=0.01%) (**Figure 1, Supplementary Table 1-2**). DN mutations were less prevalent in UKBB than LOF, and most dominant negative mutations exhibited only modest dominant negative activity (**Figure 1, Supplementary** Figure 1). We characterised an additional 2 variants in the Fenland study (see methods) which were not present in UKBB both of which we categorised as dominant negative. Our results demonstrate that damaging mutations in LXRα are prevalent in the general population.

**Figure 1.**
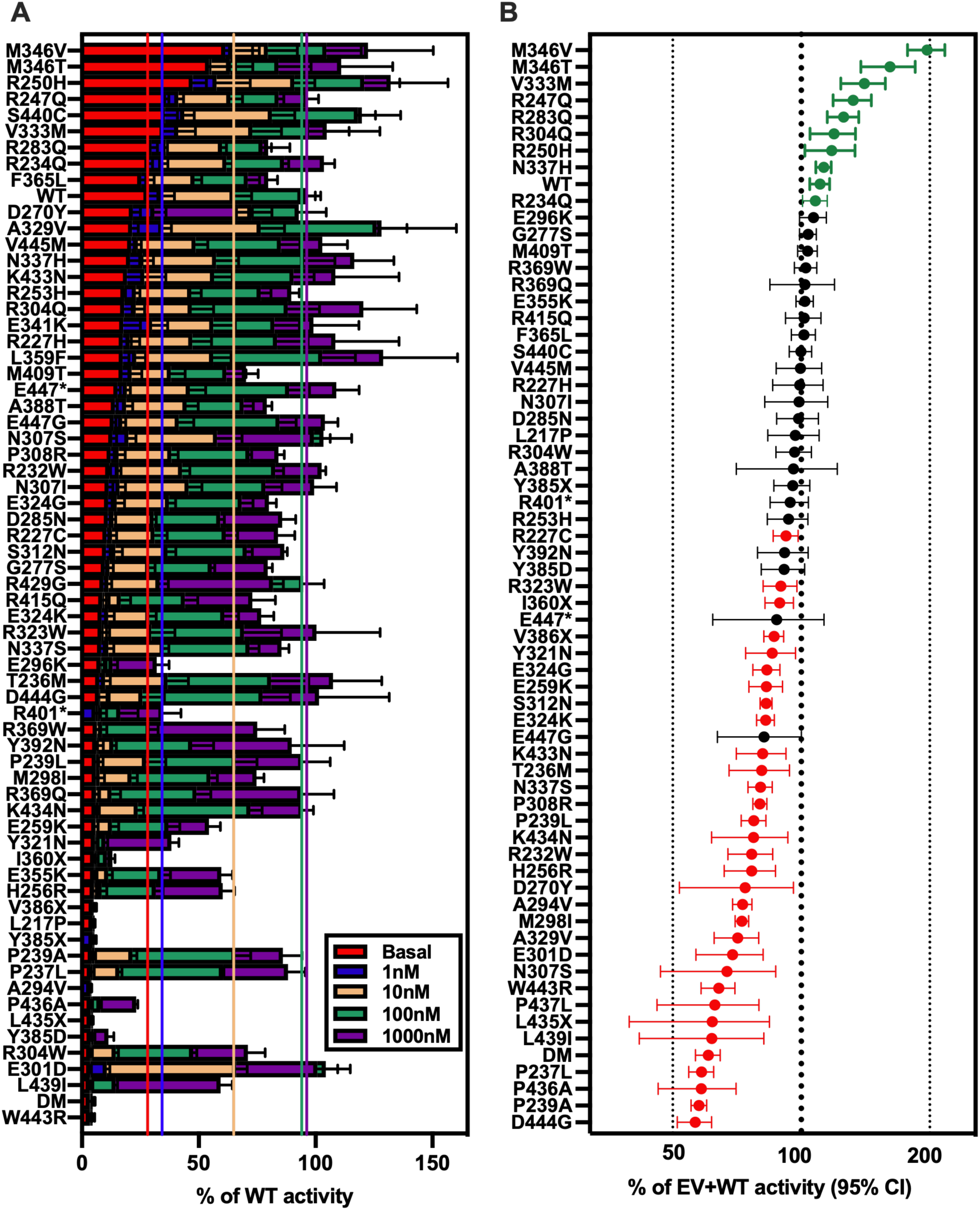
Identification of loss of function and dominant negative mutations in the ligand binding domain of LXRα carried by participants in population biobanks. We characterised 65 coding variants in the ligand binding domain of LXRα in two separate assays. A: Stacked bar chart illustrating LXRα mutant activity. HEK293 cells were transfected with an LXR-luciferase reporter construct and wildtype or mutant LXRα. 4 hours after transfection, cells were treated with indicated concentration of the LXRα agonist T0901317 and luminescence was measured as an index of LXRα activity 20 hours later. The dashed coloured lines indicate average activity of wildtype LXRα at the corresponding concentration of T0901317. Statistical analysis of mutant vs WT activity was assessed by two-way ANOVA with Geisser-Greenhouse correction and multiple comparison controlling for FDR at each ligand concentration by two-stage step-up method of Benjamini, Krieger and Yekutieli. FDR-corrected P-values are presented in Supplementary Table 1. N=3-6 experiments were performed on different day per variant in the same cell line. Height of bars and error bars represent mean +/- standard deviation. B: Forest plot of mutant LXRα activity in a co-expression assay. Wildtype and mutant LXRα and a LXR-luciferase reporter construct were transfected into HEK293 cells. 24 hours later luminescence was measured as an index of LXRα activity. Assuming normal distribution, significant increases and decreases according to a two-tailed one-sample t-test are depicted in green and red, respectively. N=4-8 independent experiments were performed on different days in the same cell line. Dots and error bars represent mean +/- 95% confidence intervals. EV= empty vector, WT= wildtype, DM = an artificial double mutant which is strongly dominant negative.

### Co-repressor association explains heterogeneity in variant effect

Nuclear receptors exhibit repressor and activator functions dependent on the repertoire of bound co-factors. Specifically, in the absence of ligand, LXRα can reside in the nucleus bound to DNA in a conformation that favours co-repressor binding thus inhibiting gene expression. Upon ligand binding, conformational changes facilitate corepressor dissociation, co-activator recruitment and receptor transactivation. Thus, disruption in co-repressor and co-activator association provide an obvious mechanistic basis for the effects of impactful variants on LXRα function.

We used mammalian two-hybrid assays to determine if differences in co-repressor or co-activator association were differentially affected by LOF and DN mutations. Both LOF and DN mutations exhibited evidence of impaired co-activator (SRC1) association in response to increasing doses of synthetic LXRα agonist **(Figure 2A)**. In contrast, co-repressor association (NCOR1) appeared relatively preserved across DN mutations but markedly impaired in all but one (p.L217P) of the LOF variants studied (**Figure 2B).** To formally compare LOF and DN mutations to WT we fitted mixed effects models with mutation class and agonist dose as interacting co-variates (see methods). Co-activator association was significantly impaired in both DN and LOF mutation groups across the majority of agonist doses tested, wheras only LOF mutations were significantly different from WT when co-repressor association was analysed **(Supplementary Table 3).** In the single gain of function mutation tested, M346V, co-activator association appeared comparable to wildtype LXRα, whereas co-repressor association was reduced. Together, these findings suggest that alteration in co-activator and co-repressor affinity of LXRα may be key determinants of variant effect.

**Figure 2.**
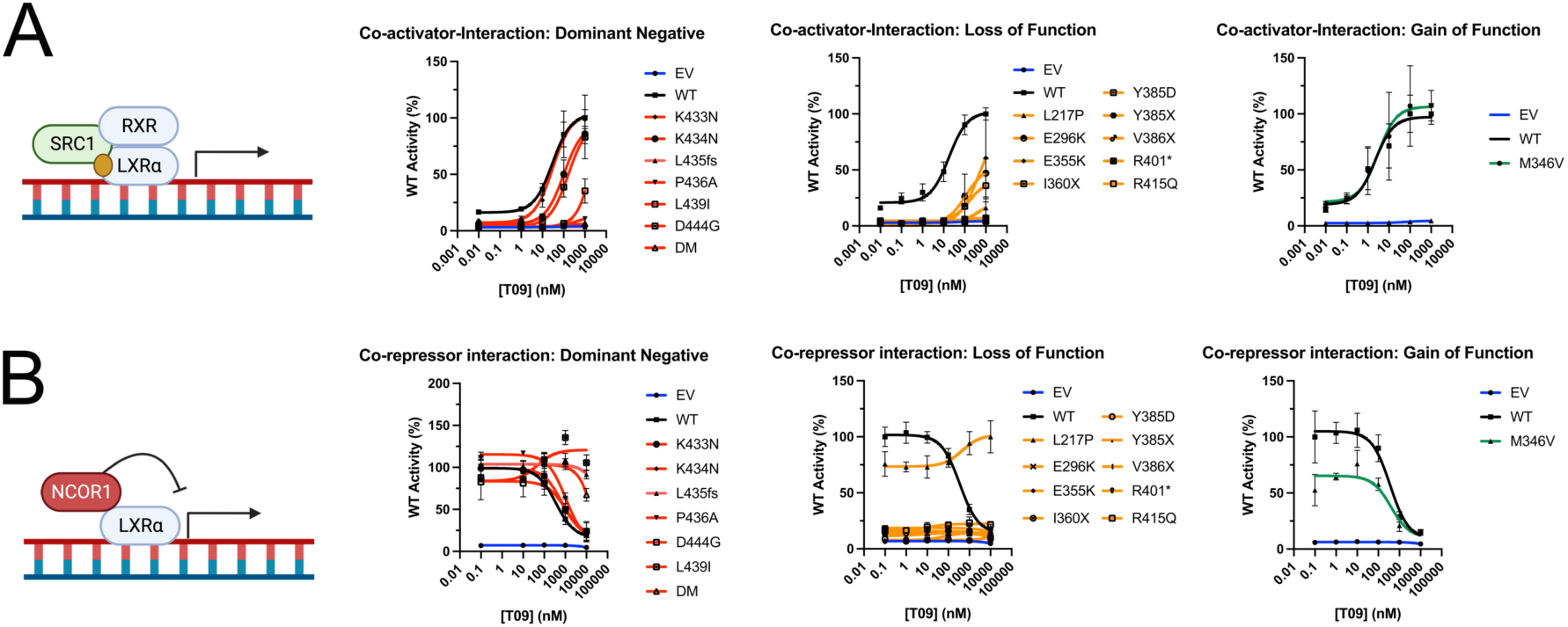
Co-repressor association is a key determinant of heterogeneity of mutant impact. We used mammalian two hybrid assays to assess the effect of mutations in the ligand binding domain of LXRα on co-activator (A) and co-repressor association (B). HEK293 cells were co-transfected with plasmids expressing LXRa-VP16, either SRC1-GAL4 (A) or NCOR1-GAL4 (B) fusion constructs with a UAStk luciferase reporter construct. 4 hours post-transfection cells were treated with the indicated concentration of T0901317 (T09) for 20 hours before luminescence was assayed as an index of VP16 and GAL4-conjugate association. Mutants were analysed in three batches each consisting of three independent experiments and are presented as normalised WT luciferase activity (%) in each batch. The mutations are separated by their classification for clarity of presentation and where possible are presented alongside the corresponding wildtype control according to experimental batch. The dominant negative and loss of function groups had mutants present in two and three batches, respectively. Therefore, the wildtype control values presented in these panels are the averages of 6 independent experiments for the dominant negative group and 9 independent experiments for the loss of function group, respectively. The curves represent lines of best fit generated in a three-parameter model conducted in Prism (Graphpad). For purposes of curve fitting and plotting on log-axis the untreated condition is plotted as an order of magnitude less than the lowest dose of agonist used. Formal hypothesis testing of mutation class on co-repressor/co-activator association versus wildtype was conducted using a mixed effects model with post-hoc dunnet’s test implemented in R4.2.2 (see methods). P-values from these analyses are presented in Supplementary Table 3. Symbols and error bars represent mean +/- SEM. EV=empty vector, WT=wildtype, DM= an artificial double mutant which is strongly dominant negative.

### Gain of function mutations in LXRα alter protein stability and co-factor affinity

To further investigate the properties of the GOF receptors we explored the ability of the receptor LBDs to interact with co-repressor and co-activator peptides in vitro. As expected from the cell-based studies, the strongest GOF mutation p.M346V exhibited two-fold reduced affinity for co-repressor peptide using a fluorescence anisotropy assay (**Supplementary** Figure 2A). Unexpectedly, the p.M346V mutant also showed a reduced affinity for co-activators in the presence of agonist. This contrasts with the cell-based two-hybrid assays which showed co-activator association for both the WT and mutant receptors (**Supplementary** Figure 2B). However, this could simply be the consequence of the co-activator concentration in the transfected cells being higher than the *Kd* of the mutant. It is notable that in cells the gain of function was most significant in the absence of added ligand (**Figure 1A**) and thus is likely to be caused by perturbation in co-repressor interaction.

The p.M346V mutation is distant from the co-repressor / co-activator recruitment surface. We therefore asked whether the mutation might mediate its effect through perturbing the global stability of the LBD. We performed thermal denaturation studies, monitored by circular dichroism, to determine melting temperatures. Both mutants p.M346V and p.M346T showed an increase in the melting temperature compared to WT and these melting temperatures don’t increase in the presence of the agonist **(Supplementary** Figure 2C). This indicates that the core of the unliganded mutant proteins is more stable than the wild-type. It is not clear how this leads to reduced co-repressor binding, but it is possible that it favours helix 12 (H12) of the receptor adopting a position that disfavours co-repressor interaction.

In **Supplementary** Figure 3 a representation of the LXRα-LBD structure and molecular modelling run by Alphafold. As mentioned above, M346 is far from the co-repressor and co-activator binding site and there are no significant structural changes predicted by Alphafold [44]. This suggests that the mutants may cause allosteric changes in the conformational equilibrium of the ligand binding domain (**Supplementary** Fig 3A-C). This agrees with the observed difference in stability. Methionine 346 is surrounded by hydrophobic residues and near two aromatic amino acids (phenylalanine 342 & phenylalanine 356) that would interact with the sulphur group and the methyl of the methionine respectively (**Supplementary** Fig 3A-C). Valine is a smaller hydrophobic residue that might preserve some of these interactions. On the other hand, threonine has a polar size chain that could potentially interact with serine 303 which in turn could also interact with serine 343. This hydrogen bond network might explain the increase in stability of the p.M346T mutant **(Supplementary** Fig 3C). Together, these findings suggest that alteration in stability of the mutant LXRα can affect co-activator and co-repressor affinity and may be a determinant of variant effect.

### Cardiometabolic consequences of damaging mutations in LXRα

To determine the effect of LXRα variants on human health, we compared carriers of experimentally proven DN, LOF, GOF and WT-like variants, as well as synonymous and bioinformatically predicted protein truncating variants (PTVs) present anywhere in the protein to non-carriers **(Supplementary** Figure 4A**, Supplementary Table 6)** in UKBB using gene burden testing. Carriers of LOF mutations that did not exhibit significant dominant negativity had higher serum HDL cholesterol, Apolipoprotein A1 and liver enzymes **(Supplementary** Figure 4A**, Supplementary Table 6)**. In general, directionally consistent findings were observed for both DN and PTV **(Supplementary** Figure 4A-B) and there was no evidence of a greater effect of dominant negative mutations across the traits tested, although the lower numbers of such mutations reduced power to detect any such additional impact and most variants exhibited only modest dominant negativity **(Supplementary** Figure 1A). When aggregated, rare (MAF<0.1%) synonymous variants - which were included as a negative control mask, did not exhibit any significant associations after adjustment for multiple testing **(Supplementary** Figure 4A**, Supplementary Table 6)**.

Given the comparability of effects across all classes of damaging mutations, we collapsed LOF, DN and PTVs to a single damaging mask to maximise statistical power (N(variants)= 70, N(carriers)= 1029, cumulative MAF=0.23%). A striking elevation of serum HDL cholesterol was observed in carriers of damaging LXRα variants (**Figure 3A, Supplementary Table 6)** and we observed a dose response relationship between LXRα variant activity and variant effect on HDL cholesterol **(Figure 3B, Supplementary Table 7)**. Damaging mutations in LXRα were associated with significant reductions in serum triglycerides consistent with loss of the known lipogenic effects of LXRα but an elevation in serum liver enzymes **(Figure 3A, Supplementary Table 6)**. These associations were robust to exclusion of the most common damaging variant, R415Q, and to adjustment for regional common variant signals **(Supplementary** Figure 4C**, Supplementary Table 8-9)**.

**Figure 3.**
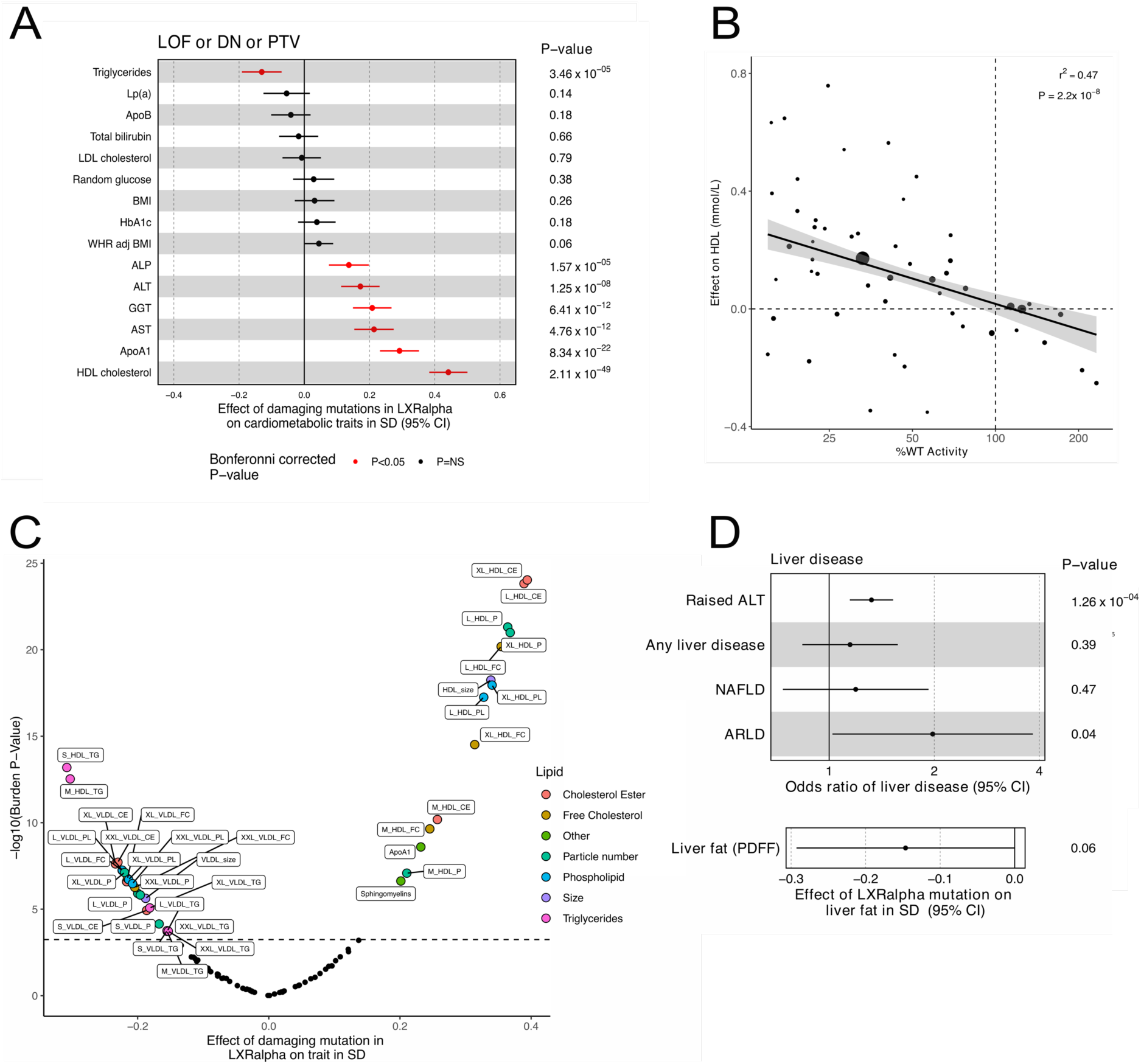
Damaging mutations in LXRα are associated with elevated liver enzymes despite beneficial effects on lipid metabolism. **A:** Forest plot illustrating the effects of carriage of a damaging mutation in LXRα (Dominant negative (DN) or Loss of function (LOF) bioinformatically predicted protein truncating variants (PTV)) on a selection of cardiometabolic traits in European participants in UK Biobank. Red points indicate statistical significance after correction for multiple testing using the Bonferonni method (P<3.9x10^-4^), NS=non-significant. **B:** Scatter plot illustrating the relationship between LXRα mutant activity and HDL cholesterol. Each dot represents an individual mutation, the size of the dot is proportional to number of carriers included in the analysis and weight in the model. The solid black line and grey band represent the estimated mean (+/- 95% confidence intervals) effect of changes in LXRα activity on HDL cholesterol derived from a weighted regression analysis. **C:** Volcano plot illustrating the effect of damaging mutations (LOF or DN or PTV) in LXRα on lipoprotein particle size and composition assessed by nuclear magnetic resonance using the Nightingale health platform in UKBB. Full list of abbreviations for lipid measurements in panel C are available in **Supplementary Table 10** The dotted line represents the Bonferroni adjusted p-value cut-off (P<5.7 x 10^-4^). **D:** Forest plot illustrating the effect of damaging mutations in LXRα on liver disease and liver fat in UK Biobank. LP(a) – Lipoprotein-a, LDL – low density lipoprotein, BMI – body mass index, WHR adj BMI – waist hip ratio adjuted for BMI, ALP – alkaline phosphatase, ALT – alanine aminotransferase, GGT – gamma glutamyl transferase, AST – aspartate aminotransferase, ApoA1 – Apolipoprotein A1, HDL – high density lipoprotein, NAFLD – non-alcoholic fatty liver disease, ARLD – alcohol related liver disease, PDFF = Proton Density Fat Fraction. P-values are burden P-values calculated in STAAR (see methods).

In view of the effects of damaging LXRα variants on serum lipids, we characterised their impact on lipoprotein composition using NMR metabolomic data available in a subset (N∼280,000) of exome-sequenced participants in UKBB. We observed elevations in cholesterol ester content in circulating HDL particles, increased HDL particle size and reduced HDL triglycerides (**Figure 3C, Supplementary Table 10-11**). This pattern is suggestive of relative CETP deficiency, an enzyme catalysing transfer of cholesterol esters from HDL to triglyceride rich lipoproteins in exchange for triglyceride and a known LXRα target gene [16]. VLDL triglyceride content was also reduced **(Figure 3C)**, consistent with the effects of LXRα on hepatic lipogenesis that has been described in pre-clinical models [1].

Damaging mutations in LXRα were associated with higher liver enzymes, suggestive of subclinical hepatotoxicity despite the known effects of LXRα on hepatic lipogenesis. We explored this relationship further and found carriers in UKBB had a 32% increased risk of clinically significant elevations in ALT and a nominal increase in risk of clinically diagnosed alcohol related liver disease but not non-alcoholic fatty liver disease, though this offservation is based on only 9 affected carriers and should be treated with caution **(Figure 3D, Supplementary Table 12)**. Liver fat measurements were only available in a small subset of UKBB, nevertheless we observed a trend to reduction in liver fat despite the elevation in serum liver enzymes **(Figure 3D Supplementary Table 12)**. Rare-variant associations with liver function tests and alcohol-related liver disease were consistent when using an independently established computational pipeline and an additional PTV-augmented model in 462,096 UKBB whole genomes of European genetic ancestry [45] **(Supplementary Table 13-16)**.

### Dominant negative mutations in LXRα cause liver injury and fibrosis in mice despite reductions in steatosis and hepatic triglycerides

Intrigued by the apparently paradoxical effects of damaging mutations in human LXRα on indices of hepatic lipogenesis and liver enzymes in humans, we wished to explore the mechanistic basis of these effects in more detail using a mouse model. We used CRISPR-Cas9 to generate a knock-in mouse model of one of the most potent dominant negative mutations we studied – LXRα W443R (W441R in mice), which was found in a single carrier in the Fenland study. This variant substitutes an aromatic residue in helix 12 of the ligand binding domain that is essential for ligand dependent activation of the receptor [46]. The AlphaFold model of p.W443R LXRα−LBD clearly shows how the change of tryptophan 443, a bulky hydrophobic residue which interacts directly with the agonist, for an arginine, a positively charged residue, changes the conformation of helix 12 displacing it from the active position (**Supplementary** Fig 5B, magenta arrows). This means that the mutation would favour the inactive conformation of H12 promoting the interaction with co-repressors and preventing the formation of the co-activator binding site. The mutant repressed LXR-regulated genes and impaired the effects of LXR agonist on LDL uptake when overexpressed in a hepatoma cell line, in keeping with dominant negativity **(Supplementary** Figure 6).

LXRα^W441R/W441R^ mice fed a low fat, high sucrose (control diet) for 8 weeks were comparable to their wildtype litter mate controls in body weight and composition at 16 weeks of age **(Supplementary** Figure 7A-C). Consistent with some features of human carriers of damaging LXRα mutations they had modest elevations in circulating ALT and AST **(Supplementary** Figure 7D-E), despite reductions in serum and liver triglycerides (**Supplementary** Figure 7F**, 7I)**. In keeping with the role of LXRα in hepatic cholesterol sensing and disposal, free cholesterol and cholesterol esters were elevated in the liver of homozygous knock-in mice (**Supplementary** Figure 7J**, 7K**). Serum HDL cholesterol was reduced (**Supplementary** Figure 7G), in line with known species-differences in HDL cholesterol metabolism between mice and humans. LXRα^W441R/W441R^ mice also exhibited evidence of increased lipid peroxidation and fibrosis, as assessed by 4-HNE and picrosirius red (PSR) staining, respectively. Inconsistent effects on immunohistochemical markers of hepatic macrophages were observed (**Supplementary** Figure 7L-R). There was no gross evidence of any extra-hepatic phenotype at necropsy, except for splenomegaly (**Supplementary** Figure 7H).

In contrast to the relatively modest phenotype observed on a low fat, low cholesterol diet, after 8 weeks of exposure to a western diet (0.2% cholesterol, 42% Kcal from fat, 34% sucrose by weight) LXRα^W441R/W441R^ mice had attenuated weight gain (**Supplementary** Figure 8A-C) and exhibited a marked elevation in circulating levels of ALT and AST despite suppression of serum and hepatic triglycerides (**Figure 4A-D, Supplementary** Figure 8D). Compared to wildtype mice, the homozygous knock-in mice exhibited a ∼8 and ∼10-fold increase in esterified and free cholesterol, respectively **(Figure 4E, Supplementary** Figure 8G). Histopathological analysis of livers demonstrated xanthogranulomatous inflammation in LXRα^W441R/W441R^ mice with immunohistochemical evidence of an allele-dependent increase in hepatic macrophages and CD3+ mononuclear cells, lipid peroxidation and hepatic fibrosis (**Figure 4F-I, Supplementary** Figure 8H-J). Extra-hepatic phenotypic assessment was notable for splenomegaly and reductions in fat pad mass **(Supplementary** Figure 8F). Histological analysis of the spleen demonstrated accumulation of lipid laden macrophages, possibly accumulating following clearance of excess tissue cholesterol (**Supplementary** Figure 8K).

**Figure 4.**
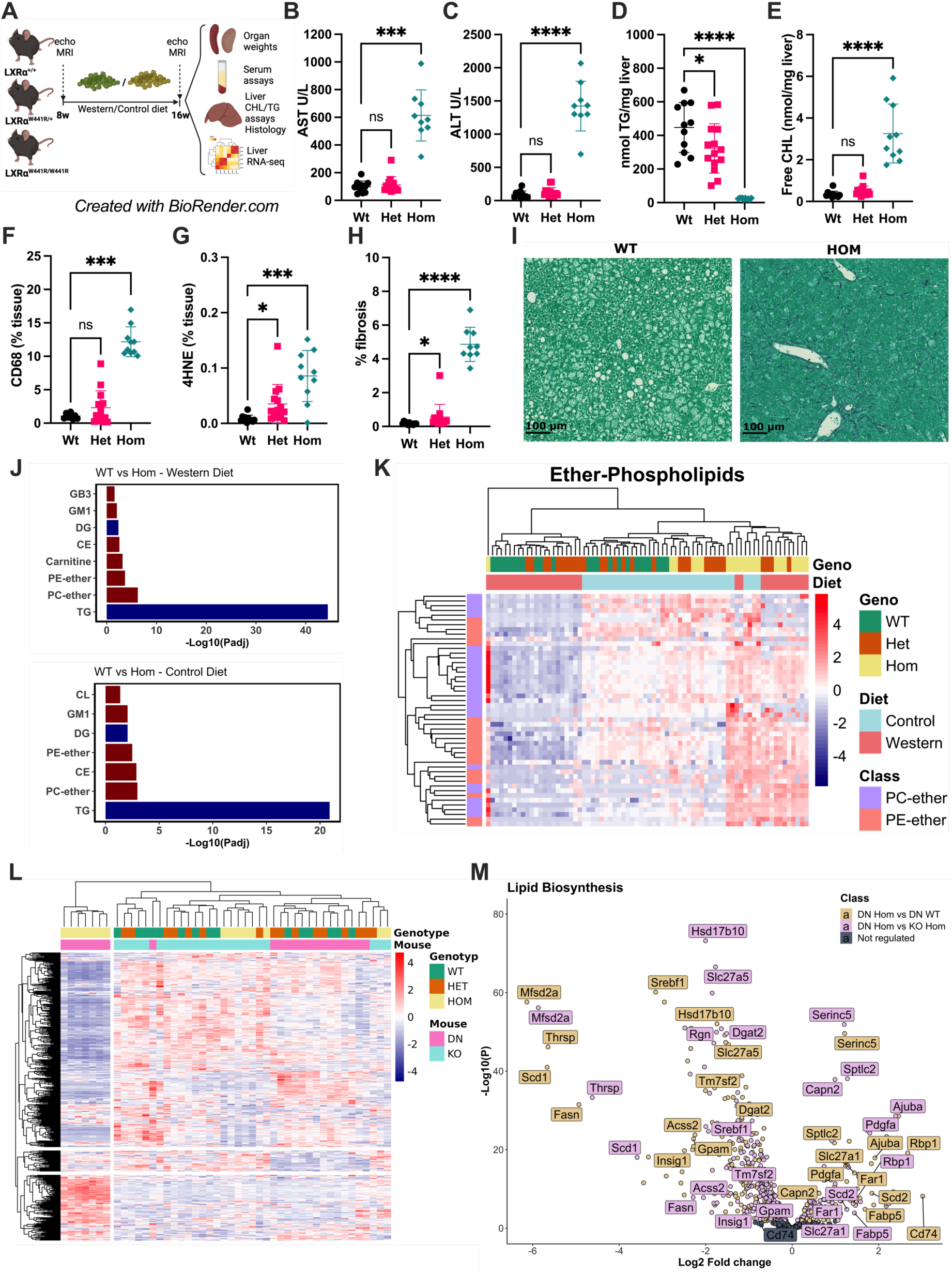
Damaging mutations in LXRα cause liver injury in mice exposed to western diet despite suppression of lipogenesis. 8-week old wildtype (LXRα^+/+^ or DN:WT, N=15), heterozygous (LXRα^+/W441R^ or DN:HET, N=19) and homozygous (LXRα^W441R/W441R^ or DN:HOM, N=14) male mice were fed Western diet (WD) for 8 weeks (**A**). After sacrifice at 16 weeks, 10 wildtype, 14 heterozygous and 10 homozygous mice were assessed for serum AST (**B**) and ALT (**C**) levels. Livers from these same mice were homogenised and assessed for level of triglycerides (**D**) and free cholesterol (**E)**. Livers were also stained for CD68 (**F**), 4HNE (**G**) and collagen using picrosirius red (PSR) stain and quantified by Halo^TM^(**H**). Representative images of PSR staining are shown in (**I**). We used mass spectrometry to assess the lipidome of Western diet and Control diet-fed mice. Enrichment of lipid classes is shown in (**J**) and a heatmap focused on Ether-Phospholipids is shown in (**K**). In a separate study, we repeated the experiment as in (**A**), with wildtype (LXRα^+/+^ or KO:WT, N=8), heterozygous (LXRα^+/-^ or KO:HET, N=8) and homozygous (LXRα^-/-^ or KO:HOM, N=8) mice. After sacrifice at 16 weeks, RNA extracted from livers of all 3 genotypes from both studies and sequenced. Heat map of RNA-seq expression data (N=8) showing sample clustering based on the genes of Lipid biosynthetic Process GO-annotation pathway (GO:0008610) (**L**), volcano plot of same genes is displayed in **M**. **B-H** were analysed by Kruskal-Wallis test with Dunn’s multiple comparison test or ordinary one-way ANOVA with Holm-Šídak multiple comparison, based on the distribution of the data. All data are presented as mean ± SD, except E which is mean ± SEM. CHL: cholesterol, MG: monoglyceride, SM: Sphingomyelin, S: Sulfatides, GB3:GB gangliosides, CL: Cardiolipin, GM1: GM1 Gangliosides, DG: Diglycerol, CE: Cholesterol Ester, PE: Phospatidylentholamine, PC: Phosphatidylcholine.

Given the key regulatory function of LXRα in liver lipid metabolism, we characterised the hepatic lipidome of our knock-in mice on a low-fat control and western diet. The most striking offservation was a marked down-regulation of hepatic triglyceride species in LXRα^W441R/W441R^ mice on control and western diet consistent with loss of the lipogenic actions of LXRα (**Figure 4J**). On both diets, there was an upregulation of ether-phospholipids, and on western diet, carnitine species increased suggesting that available fatty acid precursors may be diverted to peroxisomes and mitochondria for utilisation (**Figure 4J-K)**.

### LXRα^-/-^ mice have preserved hepatic triglyceride levels and develop less severe liver injury than LXRα^W441R/W441R^ mice

LXRα^-/-^ mice have previously been shown to develop fibrotic liver injury after prolonged exposure to a very high cholesterol diet (2% cholesterol diet for 9 months) [32]. To test if the repressive actions of the W441R variant exacerbated liver injury relative to simple loss of function we subjected LXRα^-/-^ mice to the same western diet paradigm as the knock-in mice. Unlike LXRα^W441R/W441R^ mice, weights of knockout mice were comparable to wildtype littermate controls (**Supplementary** Figure 9A-C). While LXRα^-/-^ mice exhibited some evidence of hepatotoxicity with modest rises in ALT (but not AST) and fibrosis assessed by PSR staining which appeared much more modest than the liver injury sustained in LXRα^W441R/W441R^ mice, (**Supplementary 9D-E, and Supplementary** Figure 9L-N). Free and esterified cholesterol were increased approximately 5 and 6-fold, respectively, though the change in esterified cholesterol was inflated by two mice with very high levels of esterified cholesterol (**Supplementary** Figure 9I-J). In addition, suppression of hepatic triglycerides and hepatic steatosis was not observed in knockout mice, and changes in serum triglycerides were more modest than in LXRα^W441R/W441R^ mice (**Supplementary** Figure 9F**, 9K, 9L**), suggestive of enhanced repression of lipogenic gene expression by the dominant negative p.W441R variant.

To further characterise the differential effects of LXRα knockout and carriage of a dominant negative mutation we performed transcriptomic analysis of livers from LXRα^-/-^ and LXRα^W441R/W441R^ mice. In keeping with hepatotoxic effects of loss of function in LXRα generally and its key role in lipid metabolism, we observed directionally concordant changes in inflammatory gene expression and genes related to lipid metabolism in both knockout and knockin mice relative to their littermate controls (**Supplementary** Figure 10B-D**, Supplementary Table 17-20**). However, consistent with the repressive actions of LXRα^W441R^ which were observed *in vitro*, LXRα^W441R/W441R^ mice exhibited more severe dysregulation of canonical LXRα target genes **(Supplementary** Figure 10A) and a striking downregulation in genes implicated in lipogenesis and triacylglycerol synthesis (**Figure 4L-M**) including *Fasn, Srebf1, Scd1* and *Dgat2*. We also observed striking downregulation of the Thyroid Hormone Receptor-β target gene *Thrsp* and the phospholipid transporter *Mfsd2a*. Interestingly, we observed a subset of genes implicated in fatty acid metabolism to be upregulated including the long chain fatty acid transporter *Slc27a1*, the desaturase *Scd2, Sptlc2* (sphingolipid biosynthesis) and *Far1* the rate limiting enzyme in ether lipid synthesis. Taken together these findings demonstrate, analogous to other nuclear receptors, the repressive effects of dominant negative LXRα isoforms result in more severe phenotype than loss of LXRα alone.

### Hepatocyte LXRα is hepatoprotective in mice fed a western diet

In humans and in 2 different murine models we observed uncoupling of the lipogenic actions of LXRα from hepatotoxicity. Given this apparent paradox, the fact that extra-hepatic phenotypes were observed in the LXRα^W441R/W441R^ mice and that LXRs have important immunomodulatory functions, we sought to determine if the hepatotoxic effects of LXRα^W441R^ were dependent on hepatocyte LXRα. To do this, we expressed wildtype LXRα specifically in hepatocytes using AAV8 expressing *Nr1h3* under the control of a TBG-promoter. We achieved a doubling of expression of *Nr1h3* mRNA in liver with LXRα expressing virus after 28 days (**Figure 5A**) without evidence of significant off-target expression (**Supplementary** Figure 11A-B). Consistent with the hepatoprotective effects of LXRα being exerted directly in hepatocytes, treatment with an LXRα expressing virus normalised body weight **(Supplementary** Figure 12A), prevented liver injury assessed by liver enzymes and hepatic fibrosis **(Figure 5B-C, G-H)** despite adverse effects on serum triglycerides (**Figure 5D**) and enhanced hepatic steatosis **(Figure 5G)**. Notably, adipose tissue weights were normalised, and spleen size partially normalised by hepatocyte LXRα expression, suggesting that reduced adiposity and exacerbation of splenomegaly are secondary consequences of liver injury rather than a primary effect of impaired LXR-signalling (**Supplementary** Figure 12C-E). Consistent with liver injury being dependent on hepatic cholesterol accumulation, at least in part, hepatic cholesterol levels were markedly reduced by hepatocyte LXRα expression (**Figure 5E-F**).

**Figure 5.**
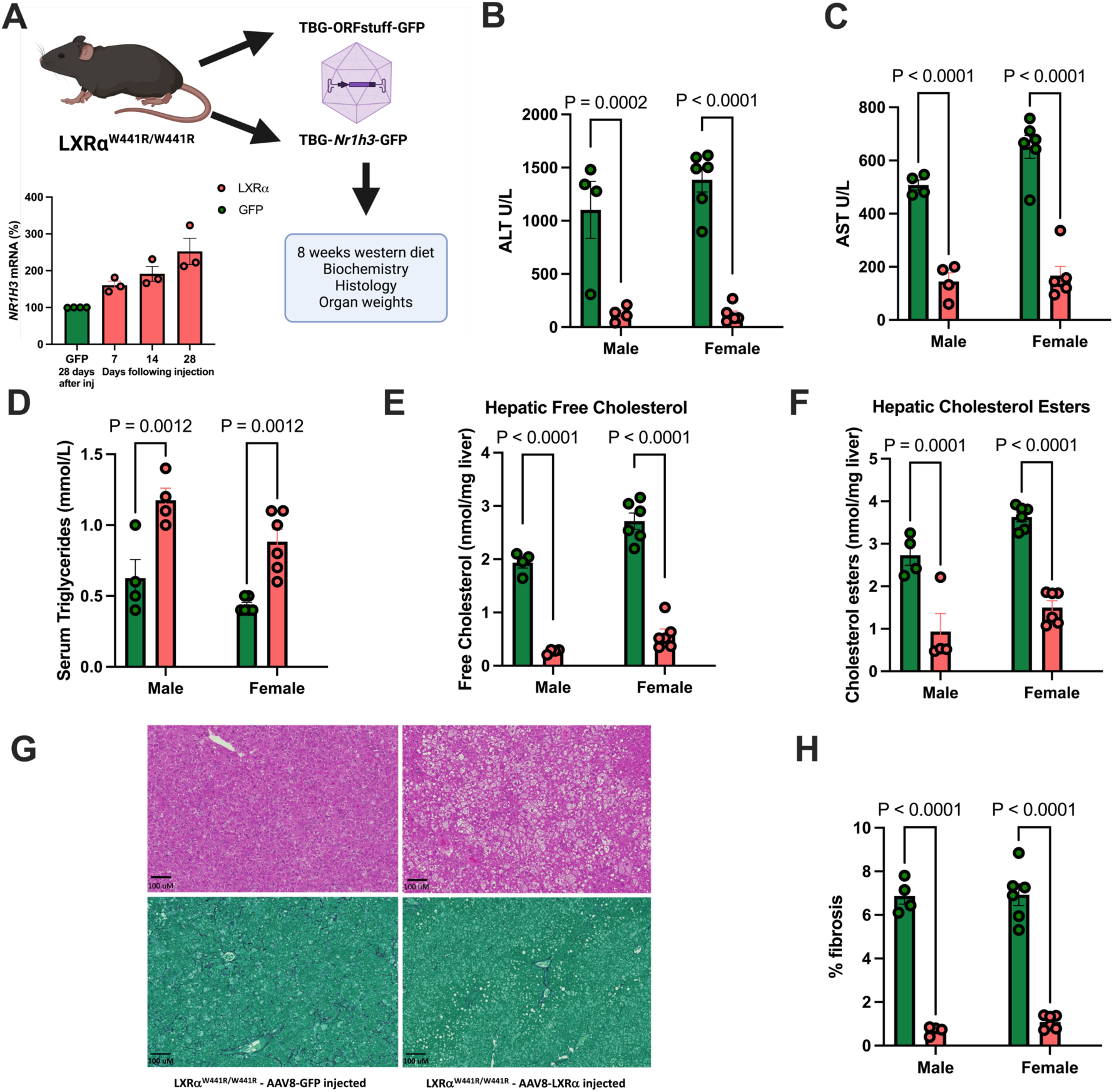
The hepatic injury that results from western diet feeding in LXRα^W441R/W441R^ mice can be rescued by hepatocyte overexpression of WT mouse LXRα. **A:** Schema of study design. 10 male and 12 female 10-12 week old C57BL/6J LXRα^W441R/W441R^ mice were randomised to receive a tail vein injection with 1x10^-11^ GC of AAV8 expressing either LXRα and GFP (LXR, salmon bars/symbols) or just GFP (GFP, green bars/symbols) under the control of the hepatocyte specific Thyroxine binding globulin (TBG) promoter (see methods) and a week later they were placed on western diet. At 4 weeks, this dose resulted in approximately two-fold induction of *Nr1h3* expression in whole liver lysate. **B-C:** Liver enzymes Alanine amino transferase (ALT) and aspartate aminotransferase (AST) after 8 weeks of western diet. **D:** serum triglycerides after 8 weeks western diet **E-F:** Free and esterified hepatic cholesterol measured using liquid chromatography with mass spectrometry detection **G:** Representative micrograph of H+E (Top) and picrosirius red (bottom) stained liver sections from each experimental group **H:** A quantification of fibrotic area derived from HALO-analysis of Picrosirius red staining (see methods). Throughout, the salmon bars and symbols represent mice treated with LXRα expressing virus and green represents mice injected with control GFP-expressing virus. All data presented are analysed with a Two-way ANOVA with post-hoc Holm-Šídak testing. Height of the bars represent mean +/- SEM.

## Discussion

Studies in cells and animal models have established that LXRα plays key roles in sensing cholesterol metabolites to regulate lipid synthesis and secretion in liver and some other tissues. However, questions remain regarding its role in human biology. By examining the impact of rare naturally-occurring, highly damaging mutations in LXR alpha on *in vivo* human phenotypes and in cells and mice expressing such mutants we have gained novel insights onto the role of this nuclear receptor in the control of human cardiometabolic and hepatic health.

Of all cardiometabolic phenotypes found in carriers of damaging mutations in LXRα studied, the strongest effect observed was on the elevation of circulating HDL cholesterol – confirming that the net effect of endogenous LXRα agonism in humans is to suppress HDL cholesterol. In contrast, synthetic LXR agonists elevate HDL cholesterol in mice [13, 15, 47]. Previous efforts to use LXR-agonism to promote reverse cholesterol transport in humans in the context of atherosclerosis were hampered, in part, by suppressive effects of LXR-agonism on HDL cholesterol [15]. It is likely that these species differences are due to the ability of LXRα to induce CETP [48], an enzyme which, in primates, exchanges triglyceride in triglyceride rich lipoproteins for cholesterol esters from HDL – thus lowering HDL cholesterol. This enzyme is absent in mice, which respond to LXR agonism with a rise in HDL [13, 15, 47]. The lipoprotein profile seen in humans carrying damaging mutations in LXRα is highly suggestive of CETP deficiency, with triglycerides carried in small and medium HDL the most downregulated and cholesterol ester in large and very large HDL the most upregulated of all lipids tested [15, 16]. Previous studies have reported a lower level of HDL cholesterol in humans with specific LOF mutations in LXRα [49, 50], Through the study of a broad range of mutations with varying degrees of functional impairment, our study reveals a dose-response relationship between LXRα transactivation capacity and circulating HDL cholesterol.

In contrast with the known effects on circulating lipids, LXRα mutations have not previously been reported to be associated with markers of hepatic injury. We found that damaging mutations in human LXRα are associated with elevations in circulating levels of liver enzymes suggestive of hepatotoxicity, despite reductions in circulating VLDL-triglycerides and a trend to reduction in liver fat. As LXRα is such a key regulator of cellular cholesterol levels in the liver [32], these findings suggest that dysregulation of hepatic cholesterol might be hepatotoxic in humans. Observational studies have provided evidence of an association between hepatic free cholesterol and presence of steatohepatitis in MASLD [51], but these studies are susceptible to confounding and reverse causation. Genetic variants which impair VLDL secretion from the liver including loss of function mutations in *APOB* and *TM6SF2* increase risk of liver disease [23] and presumably increase hepatic cholesterol, as has been shown in some rodent models [52]. However, failure to secrete VLDL particles will also cause accumulation of hepatic triglycerides, and so the effect of cholesterol accumulation due to these genetic variants cannot be dissociated from that of triglycerides. By dissociating hepatic cholesterol accumulation from hepatic lipogenesis, damaging mutations in LXRα provide a unique genetic instrument with which to interrogate the hepatotoxic effects of hepatic cholesterol accumulation in humans.

To advance our understanding of the hepatoprotective effects of LXRα we generated a knock-in mouse carrying a dominant negative mutation, p.W441R. Remarkably, these mice exhibited a striking fibrotic liver injury after 8 weeks exposure to a relatively modest dietary cholesterol challenge. Moreover, this hepatotoxicity occurred in the absence of discernible steatosis and hepatic triglycerides are markedly reduced in LXRα^W441R/W441R^ mice. While liver injury was observed in LXRα^-/-^ mice over 20 years ago [32] this occurred after several weeks on a diet with 10 times more cholesterol. Moreover, in that study, liver triglycerides were only modestly decreased after 90 days on high cholesterol diet and there was histological evidence of marked steatosis [32]. To provide a more direct comparison of LXRα^-/-^ and LXRα^W441R/W441R^ mice we undertook our own characterisation of LXRα^-/-^ on a western diet. While fibrotic liver injury was observed in knockout mice, it was much more modest in comparison to in LXRα^W441R/W441R^ mice. Moreover, hepatic triglyceride levels were comparable to wild-type. Consistent with these findings, lipogenic gene expression was suppressed to a far greater extent in LXRα^W441R/W441R^ than LXRα^-/-^ mice. Our hepatocyte specific rescue experiments confirm that the liver damage resulting from this mutation originates in the hepatocyte itself, but the precise mechanistic basis for the severe liver injury in the knock-in mice is not yet fully elucidated. It is likely that differences in hepatic cholesterol accumulation at least partly explain the more severe liver injury in LXRα^W441R/W441R^ mice: LXRα^-/-^ exhibited a 5-fold increase in free cholesterol compared to wildtype control mice, but liver free cholesterol was still ∼4-times greater in LXRα^W441R/W441R^ than LXRα^-/-^ mice. However, it is unclear if these changes are sufficient to explain the stark differences in severity of liver injury. It seems plausible that the adaptive response to hepatic cholesterol accumulation mediated by LXRα modulates sensitivity to the cytotoxic effects of cholesterol accumulation as well as mediating cholesterol disposal. For example – the lipogenic effects of LXRα could be protective in the context of cholesterol excess by providing fatty acids to esterify cholesterol in lipid droplets thereby preventing cytoplasmic cholesterol crystallisation [30]. In addition, while we have demonstrated that liver injury in LXRα^W441R/W441R^ mice can be prevented by viral expression of wildtype LXRα this does not exclude a role for impairment in LXRα in other hepatic cells in exacerbating liver injury that ensues when cholesterol accumulates due to impaired hepatocyte LXRα [53, 54]. It is conceivable that LXRβ is able to compensate for loss of LXRα in hepatocytes, but in knock-in mice the dominant negative LXRα mutant prevents sufficient compensation resulting in fibrotic liver injury. Indeed, LXRαβ−knockout exhibit enhanced fibrosis in response to non-cholesterol based hepatotoxins (CCL4 and methionine/choline-deficient diets) [53]. While the provision of a comprehensive and definitive proof of the mechanism of liver injury in the LXRα^W441R/W441R^ mice is beyond the scope of this report, these mice, with their rapid development of severe fibrosis, provide a tractable model system with which to further probe the mechanisms of cholesterol-dependent hepatotoxicity.

There are some direct translational implications of our work. Most notably, we have shown in humans and in two different mouse models that genetic impairment of LXRα is associated with hepatotoxicity. These findings caution against the use of inverse LXR-agonists, agents which are currently in clinical development for MASLD and dyslipidaemia [28]. It should be acknowledged that data characterising the effects of inverse LXR-agonists in mice have demonstrated improvements in hepatic inflammation and fibrosis in diet-induced (including high cholesterol) and alcohol-induced liver disease [26, 55–57]. However, these studies have typically been of short duration, have not demonstrated that the protective effects of these agents *in vivo* is dependent on LXR-inhibition rather than off-target effects, and have not quantitated changes in hepatic cholesterol. Unfortunately, due to the limited number of cases of liver disease in UK biobank and the rarity of damaging mutations in LXRα we were unable to conclusively demonstrate an effect of haploinsufficiency for LXRα on “hard” liver disease endpoints. We did find an increased risk of alcohol-related liver disease in carriers of damaging LXRα mutations, but this was based on a small number of affected carriers. Future studies conducted in disease cohorts of large sample size will be required to definitively assess the effect of LXRα haploinsufficiency on alcohol related liver disease and determine if LXRα modulates risk of liver cirrhosis in response to other hepatotoxic agents. Our demonstration that damaging mutations in LXRα are present in >1/500 people in the general population and are associated with evidence of sub-clinical hepatotoxicity should provide impetus for assessing the prevalence of these mutations in clinical liver disease cohorts and testing if bespoke interventions in carriers, such as very low cholesterol diets, can prevent/slow progression of their liver disease.

There are some differences between our murine and human data that are worthy of comment. While the carriage of a dominant negative mutation in LXRα was much more severely hepatotoxic in the mouse than carriage of a simple loss of function mutation, at least in the homozygous state, we did not find any evidence for greater elevation of circulating liver enzymes in humans with heterozygous dominant negative mutations compared to those with simple LOF mutations. In this regard it is worth noting that we deliberately chose a severely dominant negative mutation when generating the mouse model, while the range of dominant negativity found in the human population was wide with the majority of mutations having a considerably less adverse impact on signalling. The heterozygous knock out mice, should, a priori, be the closest model of the majority of human mutant carriers but evidence of overt liver injury was not observed in these animals. However it should be noted that the murine study was of limited duration (particularly when compared to the lifespan of human carriers in UKBB) and that C57BL/6 mice relatively poorly absorb dietary cholesterol [58] thus more severe impairments in LXRα function may be required for hepatotoxicity to manifest in these animals.

In summary, using studies in mice and humans, we demonstrate that damaging mutations in LXRα are hepatotoxic, at least in part by increasing hepatocyte cholesterol levels. Our work provides genetic evidence that intact hepatic cholesterol sensing is important in human liver health. It has also generated a novel murine model of rapid-onset liver fibrosis which should aid studies of the mechanism of cholesterol-induced hepatotoxicity.

## Methods

### Functional classification of rare variants in the ligand binding domain of LXRα

#### Variant identification and prioritisation in UK biobank

We inspected the exomes of 454,756 UK biobank participants to identify and prioritise rare variants for experimental characterisation. Processing, QC and annotation of sequencing data was undertaken as previously described, with all annotations undertaken with reference to the MANE select transcript (ENST00000441012) [59]. We selected missense variants in the ligand binding domain of LXRα with a MAF<0.001 and either a CADD score >23 or REVEL >0.7 for experimental characterisation. These are arbitrary thresholds based on the distribution of REVEL and CADD scores in damaging mutations identified in a pilot study, as well as the number of variants meeting these criteria. We also characterised a small number of protein truncating variants occurring at the C-terminus of the protein that we considered likely to act in a dominant negative manner.

#### Variant identification in the Fenland Study

To discover carriers of damaging mutations in the ligand binding domain of LXRα in the Fenland study, a population cohort study (previously described in detail [60]) we sequenced the coding region of *NR1H3* using a pooled-sequencing approach as previously described [61]. Briefly, *NR1H3* was sequenced in 11848 participants from the Fenland study using a pooled DNA approach. Equimolar amounts of DNA was pooled from 20 random participants per well at a final concentration of 10ng/ul. An amplicon DNA library was made by designing amplicons to the coding regions of *NR1H3* and PCR was undertaken using Q5® Hot Start High-Fidelity 2X Master Mix (Catalogue number M0494S, New England Biolabs® Inc) according to the manufacturer’s instructions. PCR products were purified and used in preparation of Next Generation Sequencing libraries using Nextera XT DNA Library Preparation kit (Catalogue number FC-131-1096, Illuminia®) according to the manufacturer’s instructions. Nextera XT DNA libraries were normalized with Buffer EB (Catalogue number 19086, Qiagen) to 10nM/l and each library was pooled using 5µl. Two Nextera XT DNA library pool of 376 and 216 libraries was generated for paired end 150bp, sequencing on the HiSeq4000 (Illuminia®) at the CRUK Cambridge Institute Genomics Core. Participants who carried a variant of interest needed to be identified from the DNA pool of twenty participants. This was achieved by Sanger Sequencing all of the participant DNAs from the pool individually. Variants were filtered using the criteria described above for UKBB, resulting in the identification of 2 additional variants which were experimentally characterised.

#### Cloning and site-directed mutagenesis

*NR1H3* cDNA construct (NM_005693) from pDNA3.1-hLXRα was cloned into a pcDNA3.1(+) vector (V79020, Thermo Fisher). This plasmid was used to generate mutants throughout the study. For yeast two-hybrid assays, we used pCMX-VP16-LXRα for mutant generation [62]. Site-directed mutagenesis of *NR1H3* was performed using QuikChange Lightning Site-Directed Mutagenesis Kit (210519, Agilent Technologies) according to the manufacturer’s protocols. All constructs were verified with Sanger sequencing and DNA extracted using Plasmid Maxi Kit (12163, QIAGEN). To characterize the functional consequences of LXRα mutants we performed assays in transiently transfected HEK293.

#### Cell culture

HEK293 (XX female) cells and HepG2 cells were cultured in high glucose Dulbecco’s modified eagle medium (DMEM) (41965, Thermo Fisher) and supplemented with 10% fetal bovine serum (10270, Thermo Fisher, South America origin), 1% 100X GlutaMAX (Thermo Fisher, 35050), and 100 units/mL penicillin and 100 mg/mL streptomycin (P0781, Sigma-Aldrich). Cells were incubated at 37°C in humidified air containing 5% CO_2_. Transfections were performed in HEK293 cells using Lipofectamine™ 3000 Transfection Reagent (L3000015, Thermo Fisher) and serum-free Opti-MEM I medium (31985, Thermo Fisher) according to the manufacturer’s protocols.

#### Dose-response LXRα transactivation assays

To assess the effect of LXRα variants, WT and different LXRα mutants were transiently expressed in HEK293 cells and the ligand-induced transcriptional activity measured using the Dual-Glo® Luciferase Assay System (E2940, Promega) according to manufacturer’s protocols. Briefly, 30,000 live cells were seeded in white 96-well poly-D-lysine-coated plates. After 24 hours, cells were transfected with 30 ng/well of **LXRRE luciferase plasmid** (Webster et al 1998), 30 ng/well of **pRL-TK** renilla luciferase transfection control plasmid (2241, Promega) and 40 ng/well of pcDNA3.1-LXRα or mutant plasmid. 4 hours after transfection, cell media were replaced by 75 µL of media with 0 to 1000nM of T0901317 ligand. 24 hours post-transfection, firefly and renilla luciferase activities were measured subsequently using Spark 10M microplate reader (Tecan). For normalisation, firefly values were divided by renilla and presented as %WT ± CI for each independent experiment. Each mutant was tested in a minimum of 3 independent experiments, resulting in a total of 34 independent experiments conducted across 8 individual batches. 2-way repeated measures ANOVA was computed within each experimental batch using the *afex* package, unadjusted post-hoc comparisons to the WT condition were conducted in *emmeans*, the resultant p-values were then adjusted using the Benjamini & Hochberg method to control the false discovery rate.

#### Co-expression LXRα transactivation assays for dominant negativity

Assays performed as above, with the following changes. Only 10ng/well of each of pRL-TK renilla luciferase plasmid and LXRRE luciferase plasmid were transfected per well together with 40 ng/well of pcDNA3.1-LXRα WT vector and 40 ng/well of pcDNA3.1-LXRα mutant vector. After 4 hours, media was changed only to basal (0nM T0901317) or maximum (1000nM) ligand concentration. Firefly/renilla values were normalised to % EV+WT, presented as mean ± SD, compared by one-sample t-test.

#### Mammalian two-hybrid assays

As above, cells were plated at 30,000 live cells per well in a pre-coated 96-well plate. After 4 hours, wells were transfected with 25ng of UAS-TK-Luc reporter construct, 13ng of pCMV-VP16 (EV) or pCMV-LXRα-VP16 and 13 ng of GAL4-Co-factor vector and 13ng of pRL-TK (Promega, 2241) using Lipofectamine 3000. In the case of co-activator studies, the Gal4-SRC1 (aa 570-780) [63] construct was used while Gal4-NCoR (ID 1+2) [63] was utilised for the study of co-repressor protein-protein interaction. Culture media was replaced with Fresh media containing increasing doses of the LXRα ligand T0901317 at 4 hours following transfection and incubated for a further 20 hours before Dual-Glo Luciferase Assay System. Mutants were tested in 3 separate batches with three independent experiments per batch. Results were normalised (firefly/renilla) and then presented as a percentage of average response in the 1000nM condition for co-activator (SRC1) association assays and percentage of 1nM condition for corepressor association (NCoR) within each batch. Sigmoidal dose-response curves with variable slope (three-parameter logistic regression) were fitted and plotted using Prism (Graphpad). For statistical analyses of the effects of loss of function and dominant negative mutations on co-activator and co-repressor association as an aggregated class we used mixed-effects models with normalised luciferase activity as the outcome variable and mutant class and agonist dose as fixed effects (with interactions) with individual mutant and experimental replicates included in the model as random intercepts, implemented in the Package *LmerTest*. Mutant activity was log transformed prior to analysis. As only one gain of function mutant was tested in a single batch of 3 replicates, it was assessed independently using a mixed effects model with experimental replicate included as a random intercept with normalised luciferase activity log-transformed prior to analysis. Post-hoc testing using the *emmeans* package was conducted between WT group and LOF, DN and GOF groups at each dose of agonist included in the experiment using Dunnett’s test.

#### Expression and purification of LXRα LBDs

The human LXRα WT, M346V and M346T LBDs (residues 182-447) were cloned into a pGEX2T (GE Healthcare) vector containing an amino terminal GST purification tag followed by a TEV protease cleavage site. LXRα LBDs were expressed and purified as described previously (1). Briefly, E. coli Rosetta cells were grown at 37 °C in 2xTY media and induced with 40μM isopropyl-D-1-thiogalactopyranoside (IPTG). The pellet was lysed by sonication, bound to glutathione sepharose (GE Healthcare), and washed with a buffer containing 1xPBS, 0.5% Triton X-100, 1mM DTT. Then the GST tag was removed by incubation with TEV protease (100:1 molar ratio) overnight at 4°C. Eluted proteins were loaded onto 5-ml HiTrap Q HP Ion Exchange column (HiTrap Q HP IEX), previously equilibrated in low salt buffer (20mM Tris-HCl pH 7.4, 50 mM NaCl, 1mM DTT). The protein was eluted with a 50-500nM NaCl gradient at a flow of 1.5ml/min. Pooled fractions were buffer exchanged and concentrated in 25 mM Tris/HCl pH 7.5, 100 mM NaCl and 0.5 mM TCEP buffer for fluorescence anisotropy and circular dichroism assays.

#### Fluorescence anisotropy

Co-repressor and co-activator peptides were designed for use in the fluorescence anisotropy assay. An N-terminal FITC labelled 16-aa length peptide with sequence based on the interaction domain 1 of the SMRT corepressor protein (RID1 residues 2346-2360: Ac-STNMGLEAIIRKALMG-NH2), containing the corepressor NR recognition motif LxxxIxxx[I/L]. And a N-terminal FAM labelled 16-aa length peptide with sequence based on the second NR interaction box of GRIP1 coactivator protein (NID2 residues 686-700: Ac-KHKILHRLLQDSSC-NH2) containing the coactivator NR recognition motif LxxLL.

Fluorescence anisotropy (FA) experiments were performed in black 384-well assay plates (Corning Life Sciences) as described previously [64]. Fixed concentration of SMRT and GRIP1 peptides (5nM) was used with increasing concentrations of LXRα LBDs (0-25μm) in a final volume of 20μl. For the assays in the presence of T09, increasing concentrations of the mixture protein:T09 in a 1:2 molar ratio were used. After incubation at 37 °C for 2 hours with slow shaking and centrifugation of the plates, the FA value was measured at each receptor concentration in a Victor X5 multilabel plate reader (Perkin Elmer, Singapore) using a 480-nm excitation filter and 535-nm emission filters to measure FITC and FAM emission. FA values were used to generate saturation binding curves that subsequently were used to calculate the equilibrium dissociation constant of the interaction (Kd), using the Prism software (Graphpad) and the nonlinear regression analysis.

#### Circular dichroism

Thermal unfolding of proteins was monitored by CD spectroscopy as described previously (1) using a Chirascan Spectrometer (Applied Photophysics) equipped with a temperature controller (Quantum Northwest TC125). CD spectra were measured from samples at 1mg/ml - apo - and molar ratio 1:2 (protein:agonist) when in the presence of T09 and GW.

#### LXRα LBDs structure prediction

Protein structure prediction was performed using AlphaFold2 [44], a machine-learning prediction of protein structure based on sequence and multiple sequence alignment .

#### Generation of inducible HepG2 cell lines

To generate the doxycycline-regulated lentiviral vector, human LXRα WT or W443R mutant genes were first cloned from pcDNA3.1-LXRα into the gateway-based entry vector pEN-Tmcs (ATCC-MBA-251, LGC Standard) by restriction cloning using SpeI and XhoI. Next, we cloned the LXRα gene along with the doxycycline-inducible gene cassette into the lentiviral pSLIK-Neomycin vector by site-specific recombination using Gateway™ LR Clonase™ II Enzyme mix (11791020, ThermoFisher). All clones were confirmed by restriction digest screening and sequencing. Lentiviral particles were generated as previously described [65]. Briefly, 10 μg of pSLIK-LXRα-WT or pSLIK-LXRα-W443R plasmid, 7.5 μg of each of the packaging plasmids pMDLg/pRRE and pRSREV, and 5 μg of the pseudotyping pVSV-G plasmid were co-transfected into HEK293T cells in T75 flasks using CalPhos™ Mammalian Transfection Kit (631312, Takara Bio). The culture medium was replaced 12 hours after transfection with fresh media. Viral supernatants were harvested at 24, 48, and 72 hours after transfection and concentrated by centrifugation in Centricon Plus-70 Ultracel PL-100 (UFC710008, Millipore). Lentiviral particles were added at low MOI to HepG2 cells in the presence of 8 μg/ml of polybrene (H9268, Sigma-Aldrich) for 6 hours followed by selection in 2mg/ml G418 (10131035, Sigma-Aldrich) for 3 weeks. Stable cell lines were maintained in the presence of 0.5 mg/ml G418. To avoid early activation of transgene, both selection and maintenance media contained Tetracycline-free FBS (P30-3602, Pan Biotech). Doxycycline-inducible expression of LXRα was tested by immunofluorescence, qPCR and Western Blotting.

#### Immunofluorescence microscopy

Wild-type and transgenic HepG2 cell lines were treated with 1µg/ml doxycycline for 48 hours before two PBS wash and fixation in 4% formaldehyde/PBS for 15 minutes (28906, Thermo Fisher). Cells were permeabilised with 0.1% Triton/PBS for 10 minutes and blocked for 30 minutes in filter-sterilised 3% BSA/PBS. Slides were incubated with primary antibodies for 1 hour, followed by three PBS washes and 1 hour of secondary antibody incubation. After three further PBS washes, cells were mounted with ProLong™ Gold Antifade Mountant with DNA Stain DAPI (P36935, ThermoFisher).

#### LDL-uptake assay

HepG2 cell lines transduced with inducible LXRα^WT^ and LXRα^W441R^ expressing plasmids were seeded at 200,000 per well in 24 well plates and following day treated with 1µg/ml Doxycycline. After 24 hours, cells were serum-starved with 0.1% BSA, 1µg/ml Doxycyline with or without 1uM GW3965. After 24 hours, 2.5 ug/ml Dil-LDL (Generon, LDLD15-N-1) was added. After 4 hours of incubation, cells were lysed in RIPA buffer and lysate assessed for fluorescence (Excitation: 554nm, Emission: 571) using Spark 10M microplate reader (Tecan). Signal was normalised using total protein content. Analysis was conducted by three-way ANOVA with post-hoc testing undertaken using the holm-sidak method. *P<0.05, N=4 independent experiments.

## Human Studies

### Burden testing in UK Biobank

#### Primary analysis

##### Variant masks

To understand the phenotypic effects of loss of function and dominant negative variants in LXRα in humans we performed collapsing variant tests using variant categories or ‘masks’ informed by results from experimental characterisation. We sought to identify variants in the following categories: dominant negative, loss of function without evidence of dominant negativity, gain of function, wildtype like, protein truncating variants and synonymous variants. The distribution of mutant activity in each assay was reviewed alongside number of carriers and arbitrary cut-offs selected to define each mask balancing fidelity of the functional classification and maximising carriers. These criteria are detailed in **Supplementary Table 2**. All criteria for classification were defined *a priori* before association testing. Variant lists for each mask were then used as input to the ‘collapsevariants’ applet from the MRC-EPID WES Pipeline [59] to generate a sparse matrix of variant x genotype for subsequent use in downstream association testing.

##### Phenotype preparation

For our primary analysis we pre-specified a set of 16 phenotypes (**Supplementary Table 4)** relevant to cardiometabolic health and previously described functions of LXRα from animal and cellular studies – Body mass index, WHRadjBMI, Serum biochemistry: Alanine aminotransferase, Aspartate aminotransferase, Gamma glutamyl transferase, Alkaline Phosphatase, Lipoprotein-a, Triglycerides, HDL cholesterol, LDL cholesterol, Apolipoprotein-B, Apolipoprotein-A1, Random glucose and Type 2 Diabetes. Phenotypic information was extracted from the UK biobank research access platform (UKBB RAP). Details of relevant UKBB fields and phenotype processing are described in **Supplementary Table 4**.

##### Association testing

Association testing was conducted using the MRC-EPID WES Pipeline on the UKBB RAP using the ‘extract’ function on the applet ‘mrcepid-runassociationtesting’ to test association with each variant mask against each trait of interest, as described in [59]. Briefly, this applet runs a generalised linear model (with family set to ‘gaussian’ or ‘binomial’ if trait is continuous or binary, respectively) and a STAAR [66] model. We restricted our analyses to UKBB participants of European ancestry. Age, age squared, sex, whole exome sequencing batch and the first 10 genetic principal components as defined by Bycroft et al., [67] were included as co-variates for all phenotypes. Lipid and glycaemic traits were adjusted for use of lipid lowering, blood pressure or diabetes medications as detailed in UKBB fields 6153 and 6177 (**Supplementary Table 4**).

To determine if dominant negative mutations and protein truncating variants had effects similar to ‘simple’ loss of function missense mutations we undertook a linear model regressing the effect of ‘simple’ loss of function missense mutations for each test phenotype against protein truncating and dominant negative variants using the inverse variance of the effect estimate as weights in the linear model. Finding similar effects of loss of function, dominant negative and protein truncating variants we aggregated these into a single mask of all damaging variants to maximise statistical power. We considered associations significant in our discovery analysis if the STAAR Burden P-value was less than a Bonferroni-corrected threshold of P<3.9 x 10^-4^, accounting for 128 mask x phenotype association tests.

### Sensitivity analyses

One loss of function variant, p.R415Q, is much more abundant than all other damaging variants studied (N=565) we therefore excluded this variant and repeated association testing using a variant mask including all other damaging variants, as described above. To exclude the possibility that our results were driven by known regional common variant signals, we ran genome wide association studies in UKBB participants of European ancestry for serum ALT, GGT, HDL and Triglycerides using linear-mixed models implemented in BOLT-LMM v2.3.2 [68]. Age, sex, genotyping chip and the first 10 genetic principal components were included as co-variates. We then fine-mapped the *NR1H3* (encoding LXRα) locus (11: 46790147 – 11:47790147) using Genome-wide Complex Trait Analysis (GCTA) v1.92.0 [69] to identify conditionally independent signals. We used ‘cojo-slct’ to implement a stepwise model selection procedure, defining independent associated SNPs as those with an R^2^<0.01 with a P-value of < 5 x10^-8^. The refence sample used for linkage disequilibrium estimation in the fine-mapping analysis was a random sample of 25,000 unrelated European UKBB participants. SNP dosage was extracted for the lead SNP of each conditionally independent signal for each trait using Plink v.2.00 [70] and included in a joint linear model including carrier status of a rare damaging mutation in LXRα with age, age-squared, sex, first 10 principal components, exome sequencing batch and any phenotype specific co-variates as defined in **Supplementary Table 4**.

### Follow-up analyses

#### Establishing dose response relationships between LXRα mutant activity and phenotypic effects

One attribute of experimental variant classification is the ability to test for dose response relationships between mutant function and phenotypic effects. To examine this systematically across 13 pre-specified cardiometabolic traits for activity in the transactivation assay and co-expression assay (used to assess dominant negative properties) we ran marker-level association tests in UKBB for each phenotype of interest (as defined in supplementary table 3) using BOLT-LMM v2.3.6 adjusting for age, age-squared, sex, whole exome-sequencing batch and the first 10 principal components as described elsewhere [59]. We then queried resulting summary statistics for each tested variant and regressed the effect estimate for each variant against Log_10_(mutant activity) in each assay with the inverse variance of the effect estimate used as weights in a linear model.

#### Effect of rare damaging LXRα variants on circulating NMR-derived lipid metabolites in UKBB

Given the profound effects of damaging mutations in LXRα on HDL cholesterol and its known role in lipid metabolism, we sought to examine its effects on circulating lipids in detail using the detailed nuclear magnetic resonance (NMR) spectroscopy (Nightingale Health Plc.) -based metabolomics data available in a subset of exome-sequenced UKBB participants [71]. We extracted all non-derived lipid measures (all analytes in class Apolipoproteins, lipoprotein subclasses, lipoprotein particle sizes, fatty acids and other lipids, **Supplementary Table 10**) measured in 272,281 randomly selected participants at their baseline visit. We extracted metabolites and QC flags from the UKBB cloud-based Research Analysis Platform and removed known variations during technical handling using the R package *ukbnmr* (version 2 for the phase 2 release of UKBB data) [72]. The resultant values were inverse normal rank transformed and then association testing was undertaken using the ‘Loss of function or dominant negative or protein truncating variant’ mask as described above with age, age-squared, sex, wes batch and first 10 principal components. We considered an association significant if it past a Bonferroni corrected threshold of 5.7 x 10^-4^.

#### Effect of damaging rare damaging LXRα variants on liver disease and related traits in UKBB

To assess the effect of damaging mutations in LXRα on liver health we generated liver disease and liver fat outcomes as described in **Supplementary Table 5**. Alanine aminotransferase values were defined as normal/abnormal based on clinically recommended cut-offs [72]. Liver fat measurements were derived from available abdominal MRI imaging data - Fat referenced liver proton density fat fraction (FR-PDFF) and 10-point symmetric chemical-shift encoded acquisition liver proton density fat fraction (10P-PDFF) prioritising use of FR-PDFF measurements if both were available. Liver disease outcomes were defined using self-report information, hospital episode summary and death certificate data with qualifying diagnostic codes, procedural codes and self-reported conditions outlined in **Supplementary Table 5**. ‘Any liver disease’ and controls were defined as described in Verweij et al., [23] which excluded those with raised Alanine aminotransferase levels and those with ascites without clear evidence of a non-hepatic cause from the control group. Unlike Verweij et al., [23] we included self-report in our alcohol-related liver disease classification, as well as the ICD-10 code ‘K70’. While the nomenclature and suggested diagnostic criteria for non-alcoholic fatty liver disease have recently been refined to describe a syndrome called metabolic dysfunction-associated steatotic liver disease the vast majority of cases available in UKBB will have been diagnosed before dissemination of this information and therefore a non-alcoholic fatty liver disease (NAFLD) phenotype was derived based on ICD10 codes K75.8 and K76.0. Association testing was undertaken as described above using the ‘Loss of function or dominant negative or protein truncating variant’ mask, age, age squared, sex, wes batch and the first 10 principal components were included as co-variates in all models, for liver fat the method used for liver fat measurement (10P-PDFF or FR-PDFF).

### Replication via independently established computational pipelines

#### UKBB whole genome Sequencing processing and variant calling

Whole-genome sequencing (WGS) data of the UKBB participants were generated by deCODE Genetics and the Wellcome Trust Sanger Institute as part of a public-private partnership involving AstraZeneca, Amgen, GlaxoSmithKline, Johnson & Johnson, Wellcome Trust Sanger, UK Research and Innovation, and the UKBB. The WGS sequencing methods and QC have been previously described [45, 73]. UK Biobank genomes were processed at AstraZeneca using the provided CRAM format files. A custom-built Amazon Web Services (AWS) cloud compute platform running Illumina DRAGEN Bio-IT Platform Germline Pipeline v3.7.8 was used to align the reads to the GRCh38 genome reference and to call small variants. Variants were annotated using SnpEff v4.3 [73] against Ensembl Build 38.92 [74]. We used *Peddy* and referenced 1000 genomes data [75] to classify EUR ancestry participants (peddy_prob>=0.90) removing those where peddy-derived principal components (PC) fell outside of 4 standard deviations from the mean over the first four PCs. Finally, we removed sex-discordant samples to leave 462,096 (94.2%) of unrelated EUR ancestry UKB participants for collapsing analysis.

#### AstraZeneca Centre for Genomics Research (CGR) gene-level collapsing analysis pipeline

According to the descriptions provided in previous studies [76], nine distinct models were established for collapsing non-synonymous variants, with one recessive and eight dominant models, along with an additional synonymous model as the negative control (**Supplementary Table 13**). These models were designed to consolidate functional variants that met specific criteria, referred to as qualifying variants (QVs). QVs for each model were selected based on factors such as minor allele frequency (ranging from singleton to 0.1%), predicted consequences (e.g., protein-truncating variants, missense mutations), Rare Exome Variant Ensemble Learner (REVEL) and Missense Tolerance Ratio (MTR) scores [77, 78].

Liver function test quantitative biomarkers (ALT, AST, ALP, GGT) were transformed using rank-based inverse normalisation. Four binarized clinical outcomes related to liver disease and cirrhosis (any cause, alcoholic, non-alcoholic, cirrhosis [any cause]) were referred to the definitions in Verweij [23]. Both quantitative and binarized phenotypes were subsequently regressed against individual CGR collapsing analysis model carrier status, adjusting for age at recruitment, sex, and the first four genetic principal components (PCs), using linear regression and logistic regression models, respectively. The joint PDFF phenotype underwent two independent transformations: 1) log2-transformed with outlier removal (described above); 2) rank-based inverse normalisation. Both transformed phenotypes were then regressed against the individual CGR collapsing model, adjusting for age at the corresponding release instance, sex, PDFF method (10P-PDFF or FR-PDFF), and the first four genetic PCs.

#### PTV-augmented collapsing model

A composite model was created, combining carriers of experimentally identified LXRa functional variants (**Supplementary Table 21**) with individuals carrying other PTVs independently identified in the CGR collapsing *ptv* model. This joint model was subsequently regressed against all described binary and quantitative phenotypes above, adjusting for the relevant covariates.

### Mouse studies

#### Animal housing and ethical approval

Studies were carried out at two sites: the University of Cambridge and UCLA. In Cambridge, all mouse studies were performed in accordance with UK Home Office Legislation regulated under the Animals (Scientific Procedures) Act 1986 Amendment, Regulations 2012, following ethical review by the University of Cambridge Animal Welfare and Ethical Review Body (AWERB). Mice were maintained in a 12h:12 h light:dark cycle (lights on 07:00–19:00), temperature-controlled (22 °C) facility, with ad libitum access to food and water. At UCLA, LXRα-knockout (LXRα ^−^*^/−^*) mice originally on a mixed Sv129/C57Bl/6 were obtained from D. Mangelsdorf and backcrossed more than ten generations to the C57Bl/6 background. LXRα deficient mice (Hom), heterozygous (Het), and wild type mice were maintained in a temperature-controlled room and a 12-hour light-dark cycle. Food and water were available *ad libitum*. All animal experiments were approved by the UCLA Institutional Animal Care and Research Advisory Committee. Animals at both institutes were maintained on RM3(E) Expanded Chow (Special Diets Services). Where specified, mice were fed ad libitum Western diet (TD.88137 - 42% kcal from fat, 32% sucrose by weight and 0.2% cholesterol) or Control diet (TD.05230 - 12.6% kcal from fat, 32% sucrose by weight and 0.05% cholesterol). Sample sizes were determined to detect statistically significant differences in body weight, and serum parameters between groups.

#### Generation of LXRα^W441R^ mice

Briefly, one-cell stage C57Bl/6J (Janvier-labs) embryos were injected with 50ng/ul 32fw sgRNA (IDT), 100ng/ul TriLink® CleanCap® Cas9 mRNA (L-7606) and 50ng/ul Alt-R 32fw donor top strand 123bp ssODN (IDT). Injected embryos were briefly cultured and the viable embryos were transferred the same day into pseudo-pregnant F1 (C56BL/6J/CBA) recipients [79]. F0 founder carrying the desired mutation was crossed with wild type C57BL/6J to segregate the mutations introduced and to create the F1 founder mice. Mutations were confirmed with Sanger sequencing. Founder F1 mice were crossed one more time with wild type C57BL/6J mice before establishing the mutant mouse colony. Mutations were confirmed with Sanger sequencing.

#### Genotyping strategy for LXRα^W441R^ mice

Mice were genotyped using standard PCR using primers 538_Nr1h3_amplicon_709_F1 (CCCTACAGTGGATGAGAGAGGT) and 539_Nr1h3_amplicon_709_R1 (CAACGTTAGTAGATCCCAACTGC). PCR products were digested with BglII and separated on 1.5% agarose gel. The BglII digest produces two bands (497bp and 212b) for the wildtype mice and three bands (709bp, 497bp and 212bp) for the heterozygous mice. BglII will not cut the PCR product for the homozygous mice as the mutation destroys the restriction enzyme site.

#### Mouse study 1, Western & Control diet study of LXRα^W441R^ mice

Two experimental cohorts of male homozygous (LXRα^W441R/W441R^), heterozygous (LXRα^+/W441R^) and wild-type (LXRα^+/+^) mice were generated by het × het breeding pairs. Both cohorts were weighed weekly between 4 and 16 weeks of age and had their body composition (lean and fat mass) assessed by ECHO MRI M113 mouse system (Echo Medical Systems) at 8 and 16 weeks of age. At 6 weeks of age, the first cohort of mice was transferred from standard chow to Western diet. At the same age, the second cohort was transferred to Control diet. During the experiment one mouse died unexpectedly, as such, 3 mice of each genotype of the Western Diet cohort were culled early at 14 weeks of age and pathologically assessed to exclude genotype-dependent pathology that would jeopardise animal wellfare for the remainder of the experiment. This assessment revealed reduced fat mass, splenomegaly and a hardened liver, consistent with fibrosis, as described in the main manuscript for the whole cohort, without gross evidence of other pathology. At 16 weeks of age, all remaining mice were sacrificed by CO_2_ asphyxiation and blood was isolated using cardiac puncture. Inguinal and gonadal white adipose tissues, interscapular brown adipose tissue and spleen were weighed before splitting in two, with one part fresh frozen in liquid nitrogen and the second fixed in 10% formalin. Left kidney, heart and pancreas were also isolated and fresh frozen. Blood samples were centrifuged at 4 °C and the separated plasma was stored at −70 °C until used for analysis.

#### Mouse study 2, Western diet study of LXRα^-/-^ mice

Homozygous (LXRα^-/-^), heterozygous (LXRα^+/-^) and wild-type (LXRα^+/+^) mice were generated by het × het breeding pairs. At 6 weeks of age, the first cohort of mice was transferred from standard chow to Western diet. At 16 weeks of age, the mice were weighed, and their body composition assessed by ECHO MRI (3-in-1). The mice were then sacrificed and bled by cardiac puncture. Serum, liver, inguinal and gonadal white adipose were weighed and fresh frozen in liquid nitrogen and/or fixed in 10% formalin.

#### Mouse study 3, Adenoviral rescue experiment in LXRα^W441R^ mice

To determine the dependency of the liver phenotype in LXRα^W441R/W441R^ mice, we overexpressed mouse wildtype LXRα (encoded by *Nr1h3*) using AAV8-TBG-dependent expression of *Nr1h3*. Briefly, LXRα^W441R/W441R^ mice were injected via the tail vein with 1x10^-11^ genomic copies of commercially produced (VectorBuilder) adeno-associated virus (serotype 8, AAV8) containing a vector expressing an eGFP marker gene and either, wildtype mouse LXRα (AAV8-TBG-Nr1h3) or an ORF-stuffer (AAV8-TBG-GFP), from a bi-cistronic vector downstream of the liver specific promoter TBG. After 1 week, they were placed on a Western diet and were sacrificed after 8 weeks on western diet for blood and tissue collection. Specificity of this approach was confirmed by I) Western blotting for GFP in liver (Cell Signalling, 2596S) and other tissue lysates II) Flow cytometry to identify GFP-positive cells, described below.

#### Mouse serum biochemistry

Mouse blood was collected in serum separator tubes following CO2 asphyxiation using cardiac puncture, clotted for 10 minutes at room temperature, then spun at 8,000g, serum was then collected and snap frozen on dry ice or in liquid nitrogen. Serum triglycerides, transaminases, lipoproteins and cholesterol were measured on the Dimension EXL analyser (Siemens Healthcare) or Perkin Elmer DELFIA using reagents and calibrators (Siemens).

#### Sample processing and staining for flow cytometry

Livers perfused and then collected in ice cold PBS. Tissue was then cut into small pieces and mechanically dissociated using the FFX TissueGrinder using the liver protocol. The cell suspension was then passed through a 70 μm cell strainer and RBC lysed. Cells were then stained with LIVE/DEAD Fixable Scarlet (Invitrogen) and then Fc blocked (Biolegend). Cells were then stained with conjugated antibodies: EpCAM-PE (Biolegend, clone G8.8), CD45-BV510 (Biolegend, clone 30-F11), CD31-PE-Cy7 (Biolegend, clone 390) and Ecadherin-BUV737 (Biosciences, clone DECMA-1) at 4°C for 30 minutes. Cells were then analysed using a BD Symphony A5 flow cytometer using BD FACSDiveTM Diva software. Data were analysed using FlowJo software version 10.7.1.

#### Histology and immunohistochemistry

Tissue samples were transferred from 10% formalin/PBS to 70% ethanol and processed into paraffin. 5-μm sections were deparaffinised, rehydrated, and then either stained with picrosirius red (PSR), haematoxylin and eosin (H&E) stain or were processed for immunohistochemistry (IHC) as previously described. Briefly, endogenous peroxidase activity was blocked using a 0.6% hydrogen peroxide/methanol solution. Antigen retrieval was performed using 1mM EDTA for F4/80 and CD3 or antigen unmasking solution (H3301, Vector Laboratories) for αSMA, CD68 and 4-HNE. Primary incubations were performed with 1:100 F4/80 (D2S9R, Cell signalling technology), 1:200 CD3 (MCA1477, Bio-Rad), 1:1000 FITC conjugated αSMA (F3777 Sigma), 1:200 CD68 (OABB00472 Aviva Systems Biology) and 1:200 4-HNE (Abcam, ab46545). Blocking was then performed using an Avidin/Biotin Blocking Kit (SP-2001 Vector Laboratories) followed by 20% swine serum in PBS. Sections were then incubated with primary antibodies diluted in 20% swine serum overnight at 4°C, then abiotinylated goat anti-rabbit 1:400 (Vector Laboratories), biotinylated goat anti-fluorescein1:300 (BA-0601 Vector) or goat anti-rat 1:200 (STAR80B Serotec) and then incubated with Vectastain Elite ABC reagent (PK-7100 Vector Laboratories). Staining was visualised using DAB peroxidase substrate kit (SK-4100 Vector Laboratories), then counterstained with Mayers haematoxylin prior to mounting.

H&E, PSR and immuno-stained slides were scanned (Microscopy Zeiss Axioscan Z1 Slidescanner) and processed for collagen/fibrosis (Sirius staining in % area excluding vessels) and IHC (IHC staining in % area excluding vessels) quantification using HALO image analysis software (Indica Labs).

#### RNA extraction

As fibrotic livers were not properly homogenised using ceramic beads, homogenisation was performed on all liver samples using metal beads, Trizol or Direct-zol (Zymo) and VelociRuptor V2 Microtube Homogeniser (SLS Lab Pro). Total RNAs was extracted from the mouse livers or HepG2 cells with RNeasy kit (QIAGEN) or Direct-zol RNA Miniprep Kit (Zymo Research), respectively.

#### Quantitative reverse transcription PCR

For quantitative qRT-PCR analysis, 0.5 - 1 μg of HepG2 RNA was reverse transcribed using the M-MLV Reverse Transcriptase (Promega) according to manufacturer’s protocol. qPCR reactions were prepared using TaqMan™ probes and TaqMan™ Universal PCR Master Mix (ThermoFisher) according to the protocol and were quantified using the default TaqMAN program on ABI QuantStudio 5.

#### RNA sequencing and bioinformatic analyses

RNA quality was confirmed using the Agilent 2100 Bioanalyzer or Agilent TapeStation system. For HepG2 cells, RNA-Seq libraries were prepared using the TruSeq RNA CD Indexes and the Illumina Standard mRNA Prep kit according to Illumina protocol. Multiplexed libraries were validated using the Agilent TapeStation system, normalised and pooled for sequencing. High-throughput sequencing was performed on NovaSeq 6000 (Illumina) at PE50 at CRUK Cambridge Institute Genomics Core Facility. Image analysis and base calling were performed with Illumina CASAVA-1.8.2. For mouse livers, RNA was processed by Novogene Co. Ltd. Briefly, stranded mRNA libraries were constructed by using Novogene NGS Stranded RNA Library Prep Set (PT044). The library was validated with Qubit and real-time PCR for quantification and bioanalyzer for size distribution detection. Quantified libraries were pooled and sequenced on Illumina platforms, according to effective library concentration and data amount.

Short read sequences were mapped either to human GRCh38.104 or mouse GRCm39.104 reference sequence using the RNA-Seq aligner STAR (v 2.5.0a) [80]. Differential gene expression analysis, statistical testing and annotation were performed in RStudio using DESeq2 [81]. Gene ontology and pathway analysis was performed using ShinyGO v0.77 [82].

#### Western Blot

HepG2 cells were washed in ice-cold PBS and lysed on ice in RIPA buffer (R0278, Sigma-Aldrich) supplemented with protease (1873580001, Roche/Merck) and phosphatase inhibitors (4906845001, Roche/Merck). Mouse tissue was collected at necropsy snap frozen in liquid nitrogen or dry ice and stored at -80°C before sample processing, when it was then homogenised in the RIPA-based lysis buffer described above using a FastPREP 24 Classic tissue homogeniser. Extracts were cleared by centrifufation and concentration of protein determined using Bio-Rad DC™ Protein Assay Kit (5000116, Bio-Rad). Proteins were mixed to 1X with NuPAGE™ LDS Sample Buffer (4X) (NP0007, ThermoFisher) and NuPAGE™ Sample Reducing Agent (10X) (NP0004, ThermoFisher) and denatured for 5 minutes at 95 °C. Proteins were resolved by SDS-PAGE and transferred to nitrocellulose membranes using iBlot™ 2 Transfer Stacks (Invitrogen). Membranes were blocked in 5% (w/v) skimmed milk before applying antibodies. Membranes were developed using Immobilon Western Chemiluminescent Substrate (WBKLS0500, Millipore) and imaged by ChemiDoc™ MP Imaging System (Bio-Rad).

### Lipidomics analysis

#### Liver lipid extraction and quantitation

The protein-precipitation liquid extraction protocol described previously [83]. Briefly, 15-30 mg of liver tissue was homogenised in 650 µL of chloroform with single 5 mm stainless steel ball bearing in a VelociRuptor V2 Microtube Homogeniser (Scientific Laboratory Supplies). Then 100 µL of the lipid internal standard (1-10 µM in methanol), 100 µL of the carnitine internal standard (5 µM in methanol) and 150 µL of methanol was added to each sample, followed by thorough mixing. Then, 400 µL of acetone was added to each sample. The samples were vortexed and centrifuged for 10 minutes at ∼20,000 g to pellet any insoluble material. The supernatant was pipetted into separate 2 mL screw cap amber-glass auto-sampler vials (Agilent Technologies). The organic extracts were dried using a Concentrator Plus system (Eppendorf) run for 60 minutes at 60°C.

The samples were reconstituted in 100 µL of 2:1:1 (propan-2-ol, acetonitrile and water, respectively), thoroughly vortexed and quantified by liquid chromatography with mass spectrometry detection (LC-MS) analysis, separately for cholesterol and cholesteryl-esters or full lipid profiles. The full chromatographic separation of cholesterol and cholesteryl-ester lipids was achieved using Waters Acquity H-Class HPLC System (Waters) with the injection of 5 µL onto a Waters Acquity Premier UPLC® CSH C18 column; 1.7 µm, I.D. 2.1 mm X 50 mm, maintained at 55°C. Mobile phase A was 6:4, acetonitrile and water with 10 mM ammonium formate and 0.1% formic acid. Mobile phase B was 9:1, propan-2-ol and acetonitrile with 10 mM ammonium formate and 0.1% formic acid. For the analysis of the lipid profile, formic acid was excluded from the mobile phase as previously described [83]. The flow was maintained at 500 µL per minute through the following gradient: At startL 40% mobile phase B; at 1.5 minutes: 40% mobile phase B; at 8 minutes: 99% mobile phase B; at 10 minutes: 99% mobile phase B; at 10.10 minutes 40% mobile phase B; at 12 minutes, 40% mobile phase B. The sample injection needle was washed using 9:1, propan-2-ol and acetonitrile [strong wash] and 2:1:1 (propan-2-ol, acetonitrile and water) [weak wash]. The mass spectrometer used was the Thermo Scientific Q-Exactive Orbitrap with a heated electrospray ionisation source (Thermo Fisher Scientific). The mass spectrometer was calibrated immediately before sample analysis using positive and negative ionisation calibration solution. Additionally, the mass spectrometer scan rate was set at 4 Hz, giving a resolution of 35,000 (at 200 m/z) with a full-scan range of m/z 120 to 1,800 in positive mode.

The analytes detected at the correct retention time within a 5 ppm window of the theoretical m/z were integrated along with the appropriate internal standard. The area ratio between the internal standard and the analyte were then exported and were converted into absolute amounts normalised to weight (e.g. nmoles/mg) following quality checks and blank correction.

#### Bioinformatic analyses of lipid profiles

Serum lipid profiles were analysed using the *LipidR* package [84, 85]. Missing or ‘0’ values were set to half the minimum non-zero value and then log transformed. Pairwise comparisons were undertaken between each genotype within each diet group to identify differentially regulated lipids using the ‘de_design()’ function in *LipidR* which uses a linear model to compare groups and is based on approaches developed for gene expression data [85, 86]. Lipid set enrichment analysis (LSEA) was then undertaken on the resulting dataset. LSEA is implemented in *LipidR* and is analogous to gene set enrichment though ‘sets’ in this implementation refer to lipid classes, chain length and saturation status. Enrichment scores are derived from a pre-ranked lipid list from differential expression and statistical significance of enrichment determined by permutation testing followed by correction for multiple testing [84, 85, 87, 88]. LSEA was undertaken across all contrasts tested to maintain control of Type 1 error rate. Heatmaps with hierarchical clustering of selected lipid classes were conducted using the *pheatmap* package in R.

#### Statistical analysis of mouse studies

Unless otherwise described above, studies with only male mice (mouse study 1 and 2) were analysed using Kruskal-Wallis test with Dunn’s multiple comparison test or ordinary one-way ANOVA with Holm-Šídak multiple comparison, based on the distribution of the data. Where both females and males are present, all data presented are analysed with a two-way ANOVA with post-hoc Holm-Šídak testing. Body weight curve data were analysed by mixed effects model with post-hoc Holm-Šídak test to account for repeated measures.

## Data and Code availability statement

The UK Biobank phenotype and whole-exome sequencing data described here are publicly available to registered researchers through the UKB data access protocol. Information about registration for access to the data is available at: https://www.ukbiobank.ac.uk/enable-your-research/apply-for-access. Data for this study were obtained under Resource Application Numbers 9905. Data from the Fenland cohort can be requested by bona fide researchers for specified scientific purposes via the study website (https://www.mrc-epid.cam.ac.uk/research/studies/fenland/information-for-researchers/). Data will either be shared through an institutional data sharing agreement or arrangements will be made for analyses to be conducted remotely without the necessity for data transfer. Rare variant association testing described in this paper was conducted using the MRC-EPID WES pipelines (https://github.com/mrcepid-rap/).

## Acknowledgements

SO is supported by a Wellcome Investigator Award (WT 095515/Z/11/Z). SL was funded by a Wellcome Trust Clinical PhD Fellowship (WT 225479/Z/22). RJ, EJG, KK, YJ, JRBP and KO acknowledge funding from the Medical Research Council (Unit Programme: MC_UU_00006/2). GSY, KR, APC and BYHL are supported by the MRC Metabolic Disease Unit (MC_UU_00014/1). PT acknowledges funding from the National Institute of Health (NIH R01 DK126779 and NIH P30 DK063491) and the Leducq Foundation (19CVD04). MN acknowledges funding in the form of a British Heart Foundation Intermediate Basic Science Research Fellowship (FS/20/23/34784).

Next-generation sequencing was performed via Wellcome–MRC IMS Genomics and transcriptomics core facility supported by the MRC (MC_UU_00014/5) and the Wellcome (208363/Z/17/Z) and the Cancer Research UK Cambridge Institute Genomics Core. We would like to thank core facilities at the Institute of Metabolic Science, Metabolic Research Laboratories for their help and assistance, all supported by the MRC (MC_UU_00014/5): Histology Core for assistance in histological preparation, the tissues and cell imaging core for tissue imaging and analysis, the disease models core for assistance in animal phenotyping and the metabolomics and lipidomics core for assistance with lipidomics analyses. We thank P. Barker and K. Burling of the Cambridge NIHR Biomedical Research Centre Clinical Biochemistry Assay Laboratory for their assistance with biochemical assays.

## Contribution statement

SL, MiM and SOR conceived the study. Animal model generation was conducted by IZ. Animal work was conducted by SL, MiM, IZ, MaM, JH and DR in Cambridge, AF in UCLA and JL in Newcastle supervised by SOR, APC, PT, MN, FO and DM. BYHL devised the pooled-sequencing strategy in Fenland which was executed by KR with support and supervision from GSY, YBHL, SOR, NW, MP and CL. Cellular assays assessing variant effect were conducted by SL, MiM, KD, KD and ES, supervised by KC and SOR. BA performed assessments of receptor function on purified peptides and modelling of receptor structure, supervised by JS. SL, BYHL, KK, YZ, EG, AM, RJ and FD performed statistical genetics analyses supervised by KO and JRBP. Histological assessment was conducted by SL, MiM, JT, FO and JL. Analysis of animal tissues was conducted by SL, MiM and SSM. Lipidomics analyses were conducted by AK, MiM and BJ. ‘Omics experiments were analysed by MiM and SL with bioinformatic support from BYHL. Replication of statistical genetics findings via independently established computational pipelines was conducted by XJ, KRS and DSP. SL and SOR provided funding. SL, MiM, PT, APC, KO, JRBP and SOR wrote the manuscript. All authors reviewed the manuscript for intellectually important content.

