## supplementary figures for "Damaging mutations in LXRα uncouple lipogenesis from hepatotoxicity and implicate hepatic cholesterol sensing in human liver health"

**Supplementary Figure 1. Summary of LXRα mutant effects on receptor function. A:** Scatter plot demonstrating the relationship between Transactivation capacity and co-expression activity for characterised mutations. Each dot is an individual variant, green variants are gain of function mutations, black are wildtype like, orange are loss of function without evidence of significant dominant negativity and red dots are dominant negative mutations. Grey dots are mutations which are intermediate and we not assigned to a class for the purpose of burden testing. The ‘N’ numbers represent the number of carriers in UK biobank. **B:** Schematic illustrating the effect of characterised LXR α mutants according to their position in the ligand binding domain of LXRα. Red, green and blue bars denote loss of function, gain of function and wildtype like mutations, respectively.

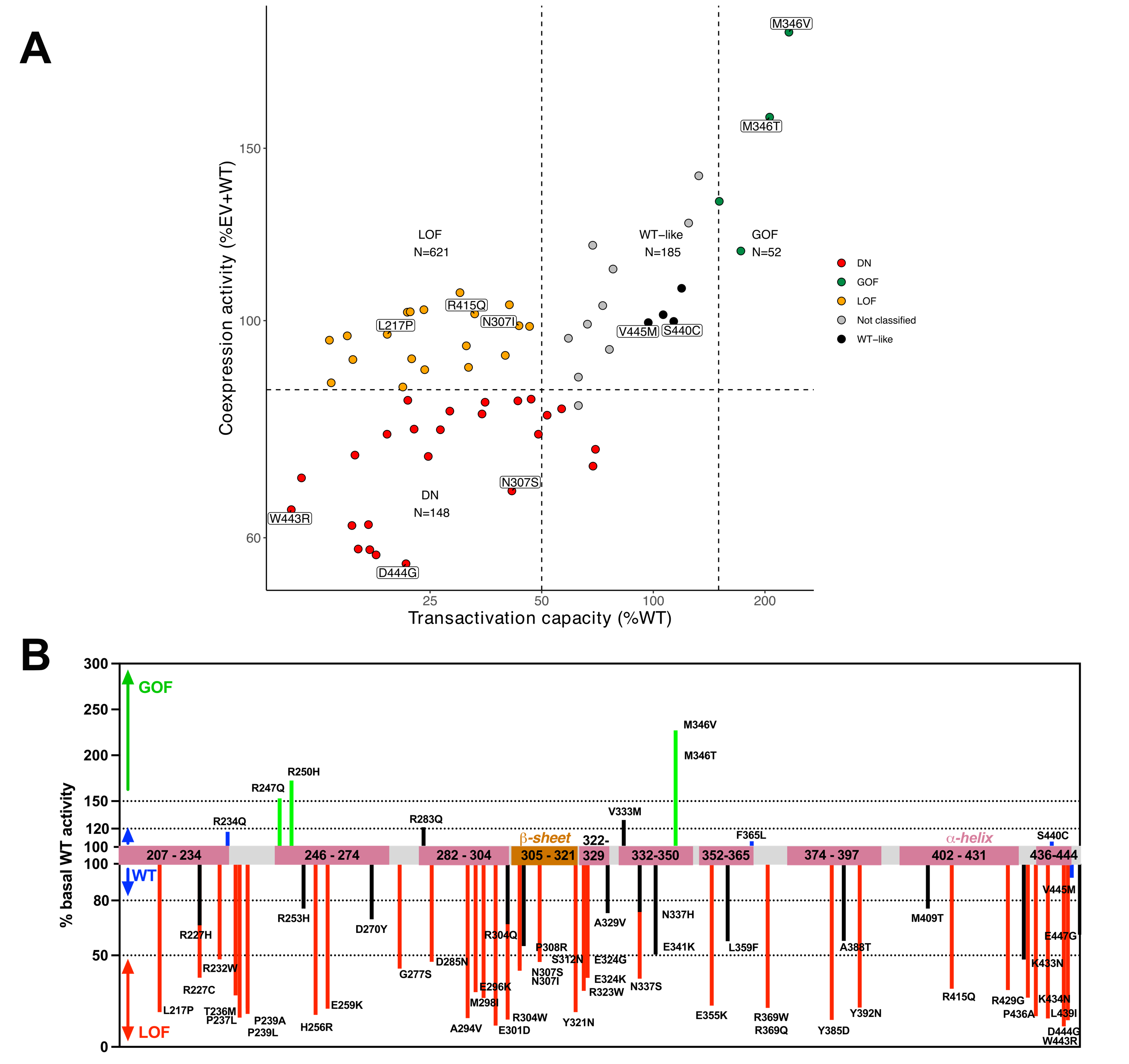

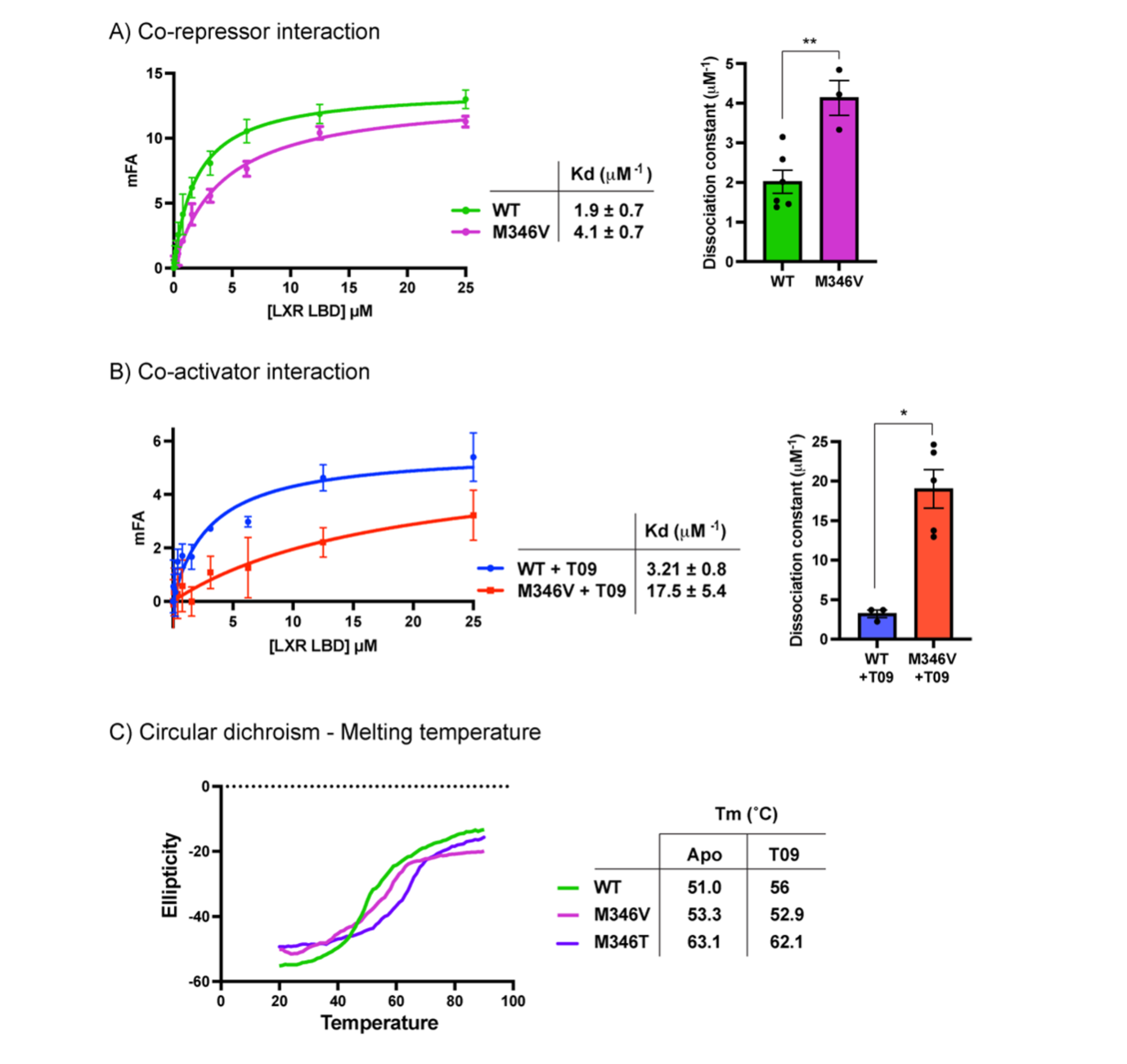

**Supplementary Figure 2. Biophysical study of gain of function mutants M346V/T LXRα by fluorescence anisotropy and circular dichroism.** Using co-repressor (A) and co-activator (B) peptides attached to a fluorophore, we studied the interaction of these peptides to the LXRα LBD mutant M346V and the thermal stability of M346V/T in vitro. A) Binding affinity in terms of dissociation constant of the WT (green) and M346V (magenta) LXRα LBDs to co-repressor peptide. B) Binding affinity in terms of dissociation constant of the WT (blue) and M346V (red) LXRα LBDs to co-activator peptide in the presence of agonist T09. C) Melting temperature of the WT (green) and M346V (magenta) M346T (purple) LXRα LBDs in the absence (apo) and presence of T09.

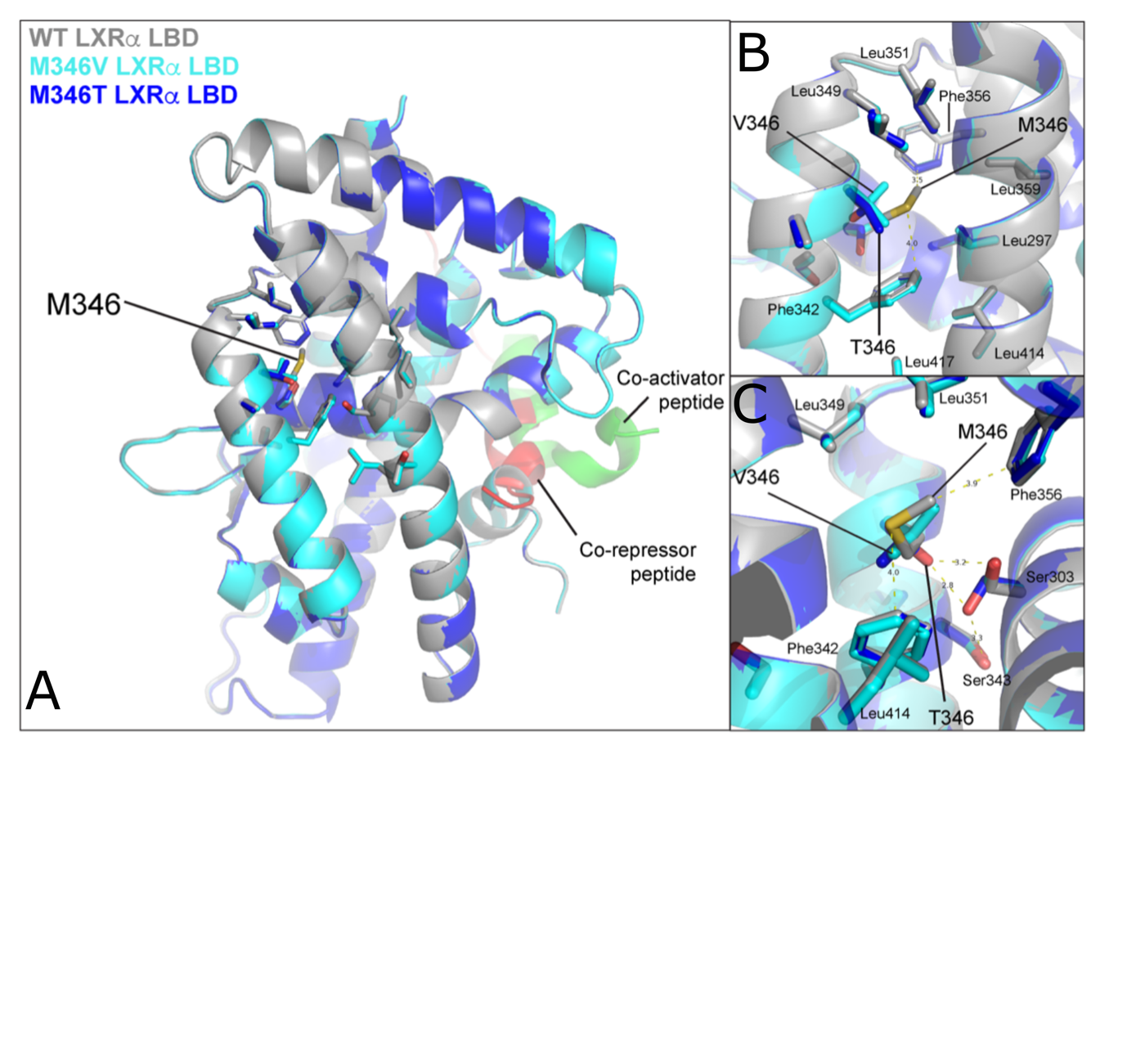

**Supplementary Figure 3. Structural models of the WT and M346V/T LXRα LBDs resulting from Alphafold. A)** Cartoon representation of the WT and M346V/T LXRα LBDs (grey, cyan and blue, respectively) models overlapping in order to highlight the differences in structure between the WT and the mutants as predicted by Alphafold. The amino acids in close proximity with the M346 (about 6 Å) are represented in sticks. The relative positions of co-repressor (red) and co-activator peptides (green) were modelled from structures PDB ID 1KKQ and 3PIQ, respectively. **B)** Close up view of the interactions between M/V/T346 and the amino acids surrounding represented as sticks. **C)** A different view of the M/V/T346 surrounding in order to highlight the potential hydrogen bonds formed by T346 with Serine 303 which would interact with Serine 343.

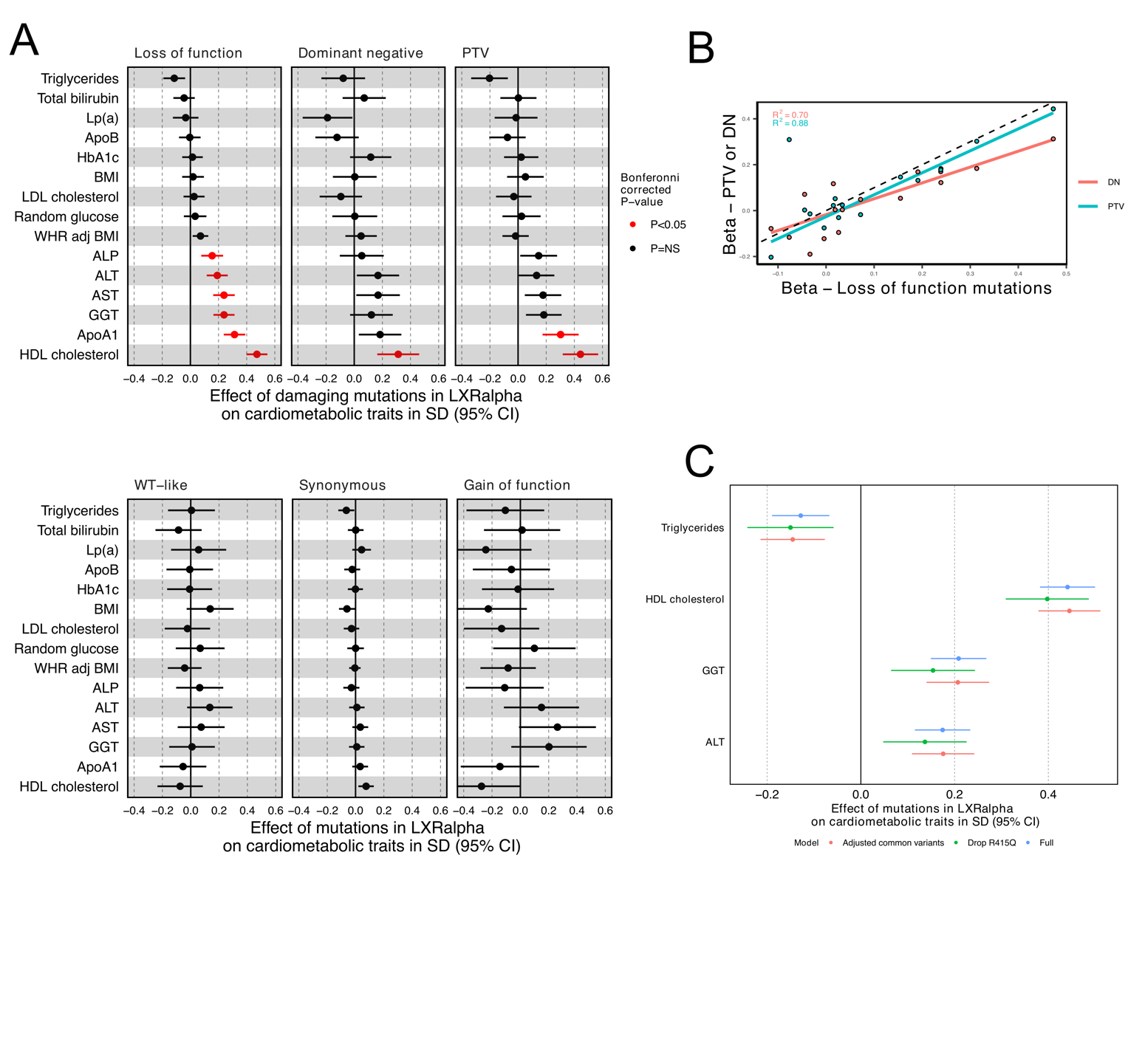

**Supplementary Figure 4. Effects of coding variants in LXRα on cardiometabolic traits. A:** The effect of coding variants in LXRα on a panel of pre-selected cardiometabolic traits. Red points indicate statistical significance after adjustment for multiple testing (Bonferroni corrected P-value – 3.9x10^-4^), NS = non-significant. **B:** Scatter plot demonstrating the relationship between the average effect of loss of function mutations and all dominant negative (DN) or protein truncating variants (PTV) on cardiometabolic trait. Each dot plots the effect of the Loss of function mutations on a single trait against either PTVs (blue) or DN (red). The dashed line represents y=x where there would be perfect agreement between the effects of Loss of function mutations and DN/PTV.  **C:** Forest plot of sensitivity analyses. ‘Full’ illustrates the effect of damaging mutations in LXRα (LOF or DN or PTV) from the primary analysis, Drop R415Q – analysis after removing the most common rare variant (R415Q, N(carriers in all UKBB) = 565) or ‘Adjusted common variants’ - adjusting for common variants in the region significantly associated with the listed traits of interest. LP(a) – Lipoprotein-a, LDL – low density lipoprotein, BMI – body mass index, WHR adj BMI – waist hip ratio adjusted for BMI, ALP – alkaline phosphatase, ALT – alanine aminotransferase, GGT – gamma glutamyl transferase, AST – aspartate aminotransferase, ApoA1 – Apolipoprotein A1, HDL – high density lipoprotein

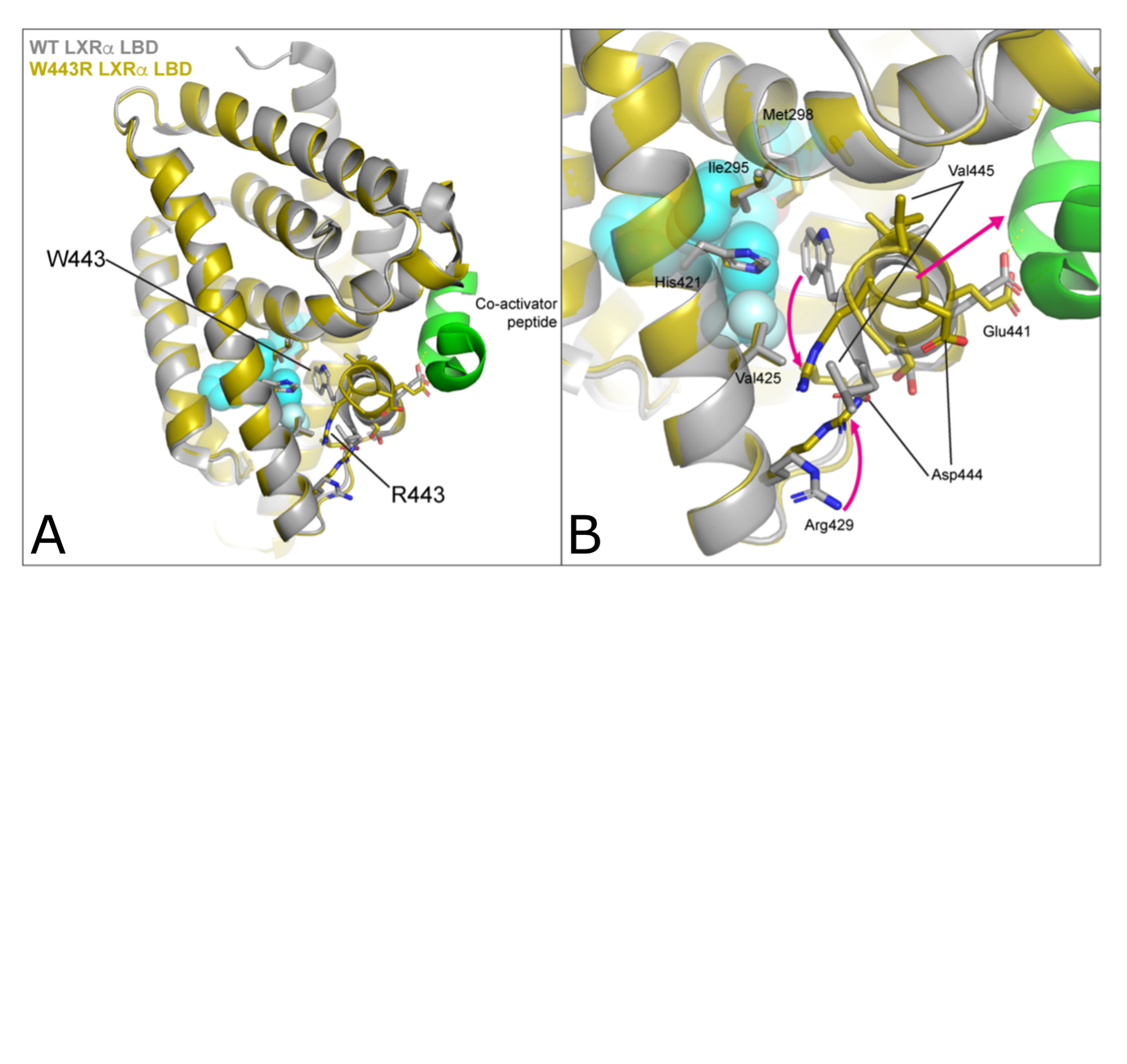

**Supplementary Figure 5. Differences between the WT and W443R LXRα LBDs Alphafold models.** Cartoon representation of the WT (grey) and W443R (yellow) LXRα LBDs models overlapping. The amino acids in close proximity with the W443 are represented in sticks. The relative positions of the agonist GW and co-activator peptide (green) were modelled from structures PDB ID 1KKQ and 3PIQ, respectively. The magenta arrows represent the relative movement of the W443 H12 position to the R443 H12 position according to the Alphafold model. Mutation W443R might potentially affect the interaction with the agonist and the active position of H12 which in turn would disturb the interaction with co-repressor and co-activator.

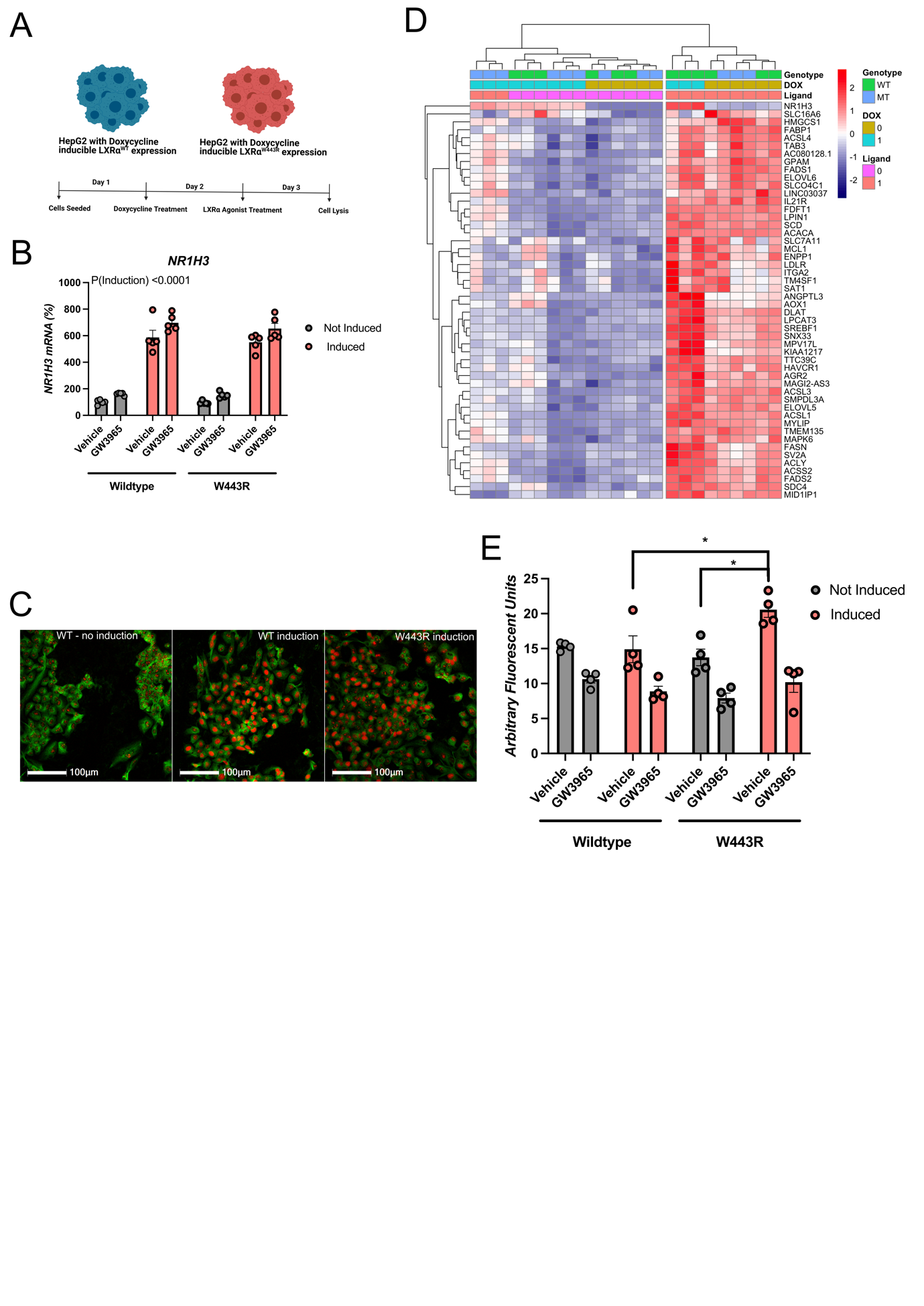

**Supplementary Figure 6 A naturally occurring variant in LXRα exerts dominant negative effects in cultured hepatocytes. A:** To demonstrate dominant negative effects of LXRα^W443R^ in a relevant cell type we generated HepG2 cell lines stably transfected with a Doxycycline-inducible transgene expressing either a wildtype LXRα isoform (WT) or a mutant LXRα^W443R^ (MT or W443R). The experimental paradigm is summarised in (A), 24 hours after seeding cells were treated with doxycycline to induce LXRα expression, 24 hours later LXR agonist GW3965 (1000nM) was added and cells were lysed 24 hours later. **B:** Expression of *NR1H3* mRNA (encoding LXRα) in the paradigm described in (A). N=5 independent experiments. P-values are displayed from 3-way ANOVA with post-hoc Holm-Šídak test. **C:** Immunofluorescent images of non-transduced HepG2 cells and WT or MT/W443R HepG2 cells treated with doxycycline stained for Beta-Tubulin, LXRα and DAPI. Enhanced LXRα nuclear immunoreactivity can be seen in the transduced cells consistent with induction of LXRα    **D:** Heatmap illustrating expression of GW3965 regulated genes in an RNAseq experiment after treatment of cells in the paradigm outlined in (A). Ligand treated cells with overexpression of LXRα^W443R^ cluster with non-ligand treated conditions, indicative of the inhibitory effects of LXRα^W443R^ on LXR-dependent signalling.  Furthermore, induction of canonical LXR-regulated genes, including *FADS1, SCD, ANGPTL3, LPCAT3, FASN, MYLIP and SREBF1,* is impaired by induction of LXRα^W443R^, consistent with dominant negative actions of LXRα^W443R^. N=3 independent experiments. **D:** We further validated the dominant negative actions of LXRα^W443R^ in hepatocytes in an LDL uptake paradigm. Following the experimental paradigm illustrated in (A), cells were treated with Dil-labelled LDL for 4 hours before cells were lysed and fluorescence measured as an index of LDL uptake. LXRα^W443R^ induction was capable of increasing LDL uptake in the basal state, counteracting the previously described effects of LXRα in this setting [1]. Analysis was conducted by three-way ANOVA with post-hoc testing undertaken using the  Holm-Šídak method. *P<0.05, N=4 independent experiments.

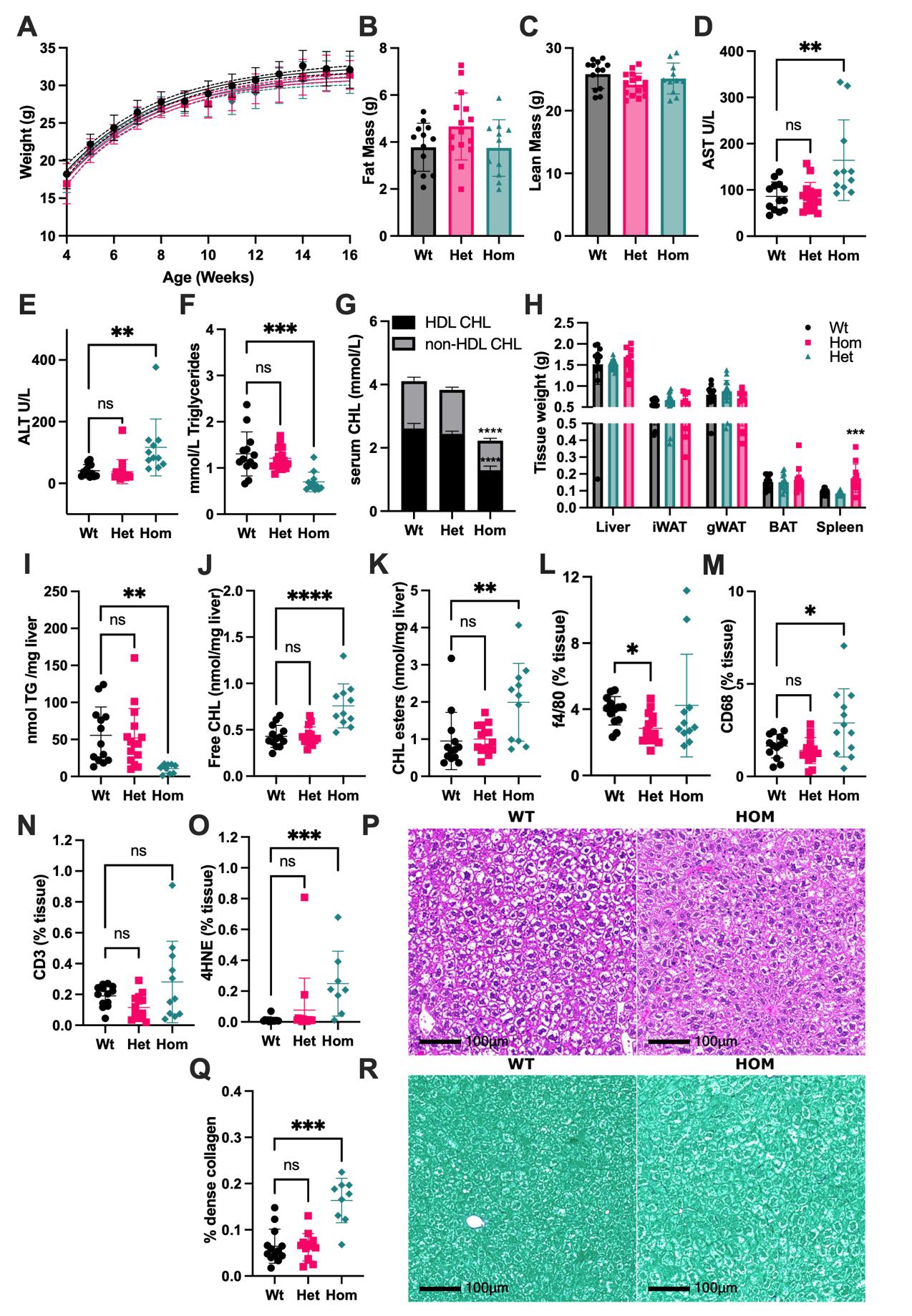

**Supplementary Figure 7. Hepatotoxicity of a dominant negative LXRα mutation in mice fed a low fat diet.** 8 week old LXRα**^WT/WT^** (Wt, N = 13), LXRα^+/W441R^ (Het, N = 15) and LXRα^W441R/W441R^ (Hom, N = 11) mice were placed on a low fat (12% by Kcal), high sucrose (34.1% by weight) diet for 8 weeks when mice were euthanised and blood and tissues were collected for downstream analyses. **A:** Body weight curves (mean and 95% confidence intervals). **B-C:** Lean and fat mass determined by Echo-MRI at 16 weeks of age. **D-G:** Serum biochemistry from terminal bleeds, in **(G)** the asterix represent significant differences between Wt and Hom mice for HDL-cholesterol and all (HDL + non-HDL) cholesterol, there was no effect of genotype on non-HDL cholesterol **H:** Tissue weights, **I-K:** lipid measurements in liver lysate normalised to liver weight. **L-O:** Immunohistochemical staining for indicated markers was performed and quantified using HALO image analysis platform (Indica labs). Representative images of H&E staining are shown in (**P**). Livers were stained for collagen using picrosirius red (PSR) stain and quantified by Halo^TM^(**Q**). Representative images of PSR staining are shown in (**R**). **A:** Body weight curves were compared by mixed effects model was used with post-hoc Holm-Šídak test. **B-O:** Analysis was performed using Kruskal-Wallis test with Dunn’s multiple comparison test or ordinary one-way ANOVA with Holm-Šídak multiple comparison, based on the distribution of the data. All data are presented as mean ± SD. *P<0.05, **P<0.01, ***P<0.001, ****P<0.0001.

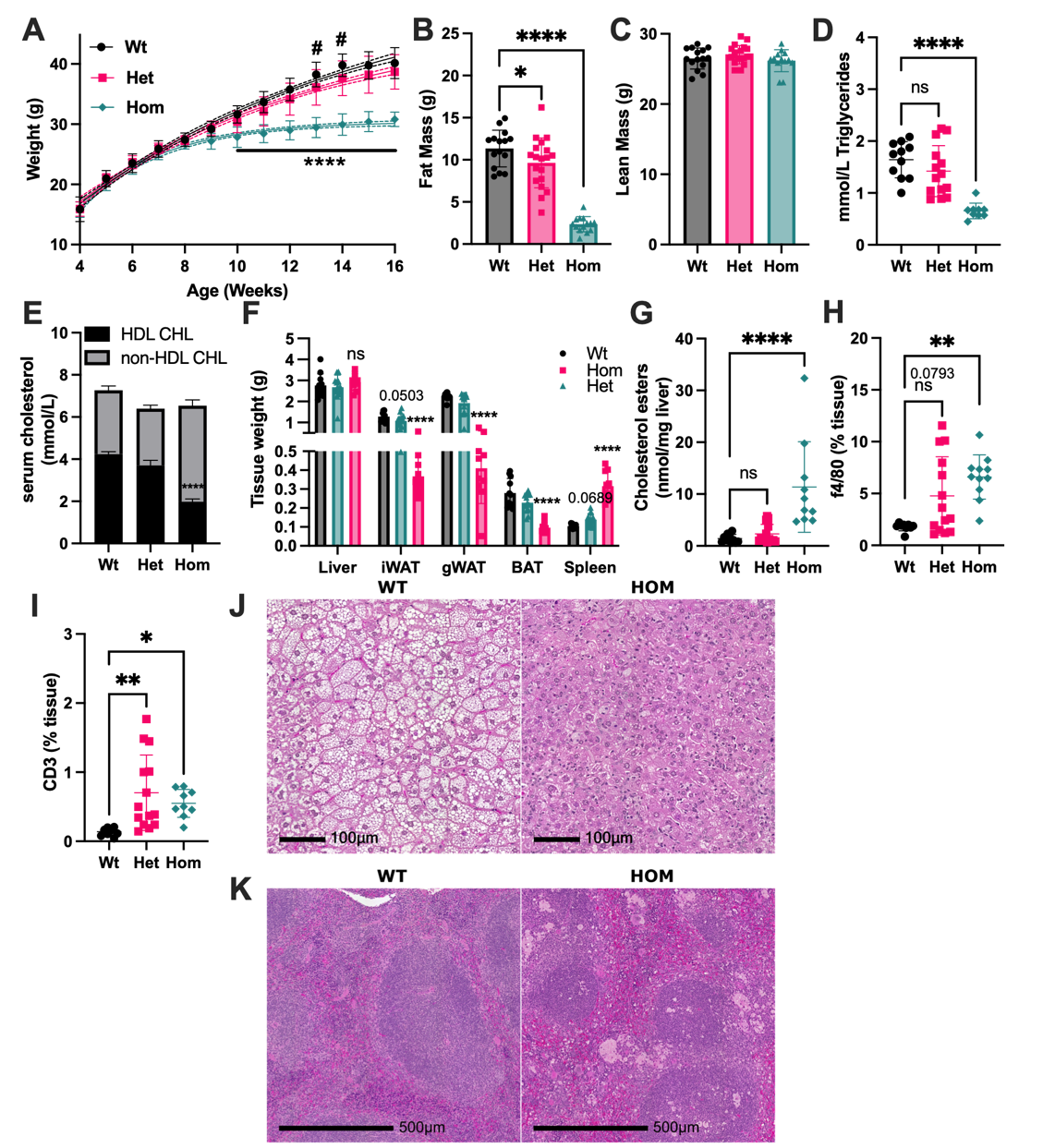

**Supplementary Figure 8. Effects of a dominant negative LXRα mutation in mice fed a western diet for 8 weeks.** 8-week-old LXRα^+/+^ (Wt, N=15), LXRα^+/W441R^ (Het, N=19) and LXRα^W441R/W441R^ (Hom, N=14) mice were placed on a high fat (42.7% by Kcal), high sucrose (34.1% by weight), high cholesterol (0.2%) diet (western diet) for 8 weeks then euthanised and blood and tissues were collected for downstream analyses. **A:** Body weight curves (mean and 95% confidence intervals). ****P<0.0001, Wt vs Hom, # P<0.05 Wt vs Het. **B-C:** Fat and lean mass determined by Echo-MRI at 16 weeks of age. For 11 wildtype, 14 heterozygous and 10 homozygous mice we performed downstream analysis including: serum biochemistry from terminal bleeds (**D-E**). In **(E)** the asterixis represents significant differences in HDL cholesterol, there was no effect of genotype on non-HDL cholesterol. **F:** Tissue weights **G-I:** Immunohistochemical staining for indicated markers was performed and quantified using HALO image analysis platform (Indica labs). Representative images of H&E staining of liver and spleen are shown in **J** and **K**, respectively. In **(A)** a mixed effects model was used to account for repeated measures with post-hoc Holm-Šídak test. In (**B-H)**, analysis was performed using Kruskal-Wallis test with Dunn’s multiple comparison test or ordinary one-way ANOVA with Holm-Šídak multiple comparison, based on the distribution of the data. Data are presented as mean ± SD. *P<0.05, **P<0.01, ***P<0.001, ****P<0.0001.

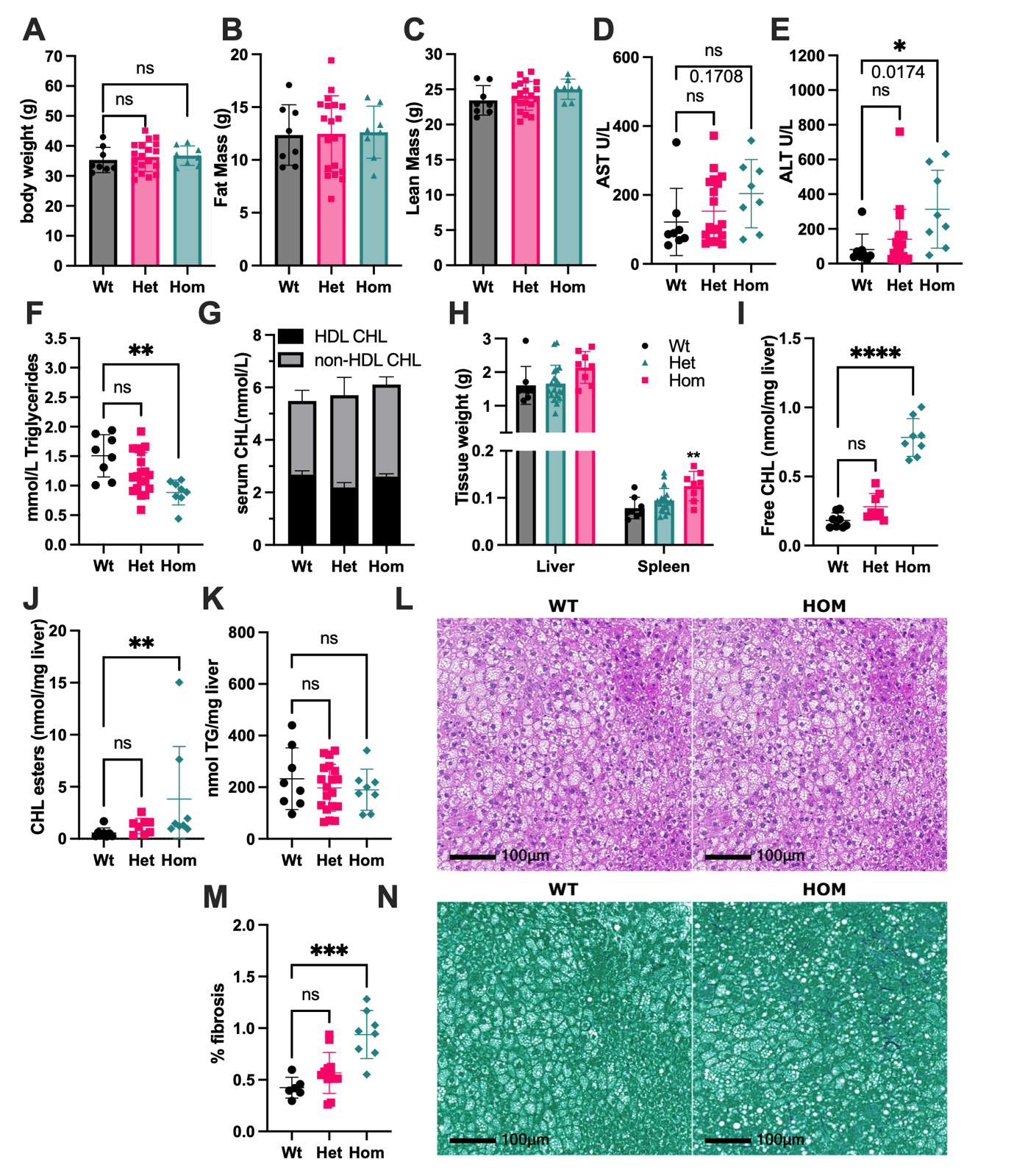

**Supplementary Figure 9. Hepatotoxicity in LXRα knockout mice fed a western diet.** 8-week-old wildtype LXRα^+/+^ (Wt, N=8), heterozygous LXRα^+/-^ (Het, N=19) and homozygous LXRα^-/-^ (Hom, N=8) mice were placed on a high fat (42.7% by Kcal), high sucrose (34.1% by weight), high cholesterol (0.2%) diet (western diet) for 8 weeks then euthanised and blood and tissues were collected for downstream analyses. Unless specified, all mice were assessed for: **A**: Endpoint weight measurements **B-C:** Fat and lean mass assessed by Echo-MRI at 16 weeks of age. **D-G**: Serum biochemistry from terminal bleeds. **H:** Selected organ weights. **I-J:** Free cholesterol (CHL) and CHL ester measurements in liver lysate normalised to organ weight are shown for N=8 mice of each genotype. **K:** Triglyceride (TG) measurements in liver lysate normalised to organ weight are shown for all mice. Representative images of H&E staining are shown in (**L**). Livers from wildtype (N=8), heterozygous (N=12) and homozygous (N=8) mice were stained for collagen using picrosirius red (PSR) stain and quantified by Halo^TM^(**M**). Representative images of PSR staining are shown in (**N**). In (**A-K, M)**, analysis was performed using Kruskal-Wallis test with Dunn’s multiple comparison test or ordinary one-way ANOVA with Holm-Šídak multiple comparison, based on the distribution of the data. All data are presented as mean ± SD. *P<0.05, **P<0.01, ***P<0.001, ****P<0.0001.

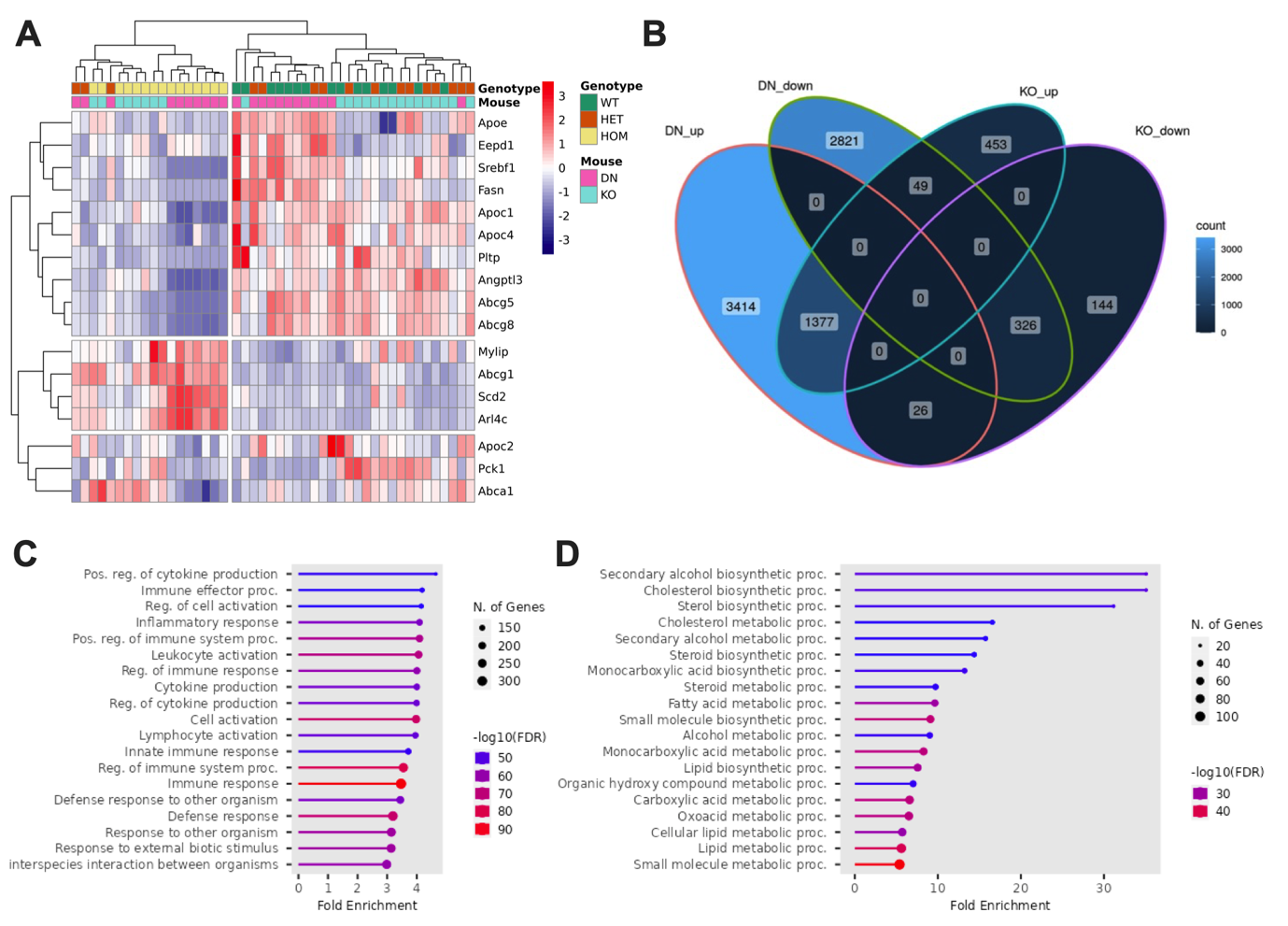

**Supplementary Figure 10.** Liver RNA-seq analysis of male mouse livers at 16 weeks of age after 8 weeks on Western Diet. 8 wildtype (LXR⍺^+/+^), 8 heterozygous (LXR⍺^W441R/W441R^) and 8 homozygous (LXR⍺^W441R/W441R^) mice of the LXR⍺-W441R dominant-negative line (DN) were used. For the LXR⍺-KO (KO) mouse model, 8 wildtype (LXR⍺^+/+^), 8 heterozygous (LXR⍺^+/-^) and 8 homozygous (LXR⍺^-/-^) mice were used**.** Heat map of RNA-seq expression data showing sample clustering based on mouse genes homologous to Reactome genes in ‘*NR1H2 and NR1H3-mediated signalling*’ (R-HSA-9024446) (**A**). Venn diagram showing overlap of up and down-regulated genes in HOM vs WT in both DN and KO mouse lines (**B**). Gene Ontology (GO) pathway analysis of 1377 shared upregulated (**C**) and 326 downregulated genes (**D**) in both (LXRα^W441R/W441R^ and LXRα^-/-^) homozygous mouse models highlighting upregulation of inflammatory genes and downregulation of genes related to sterol and fatty acid metabolism.

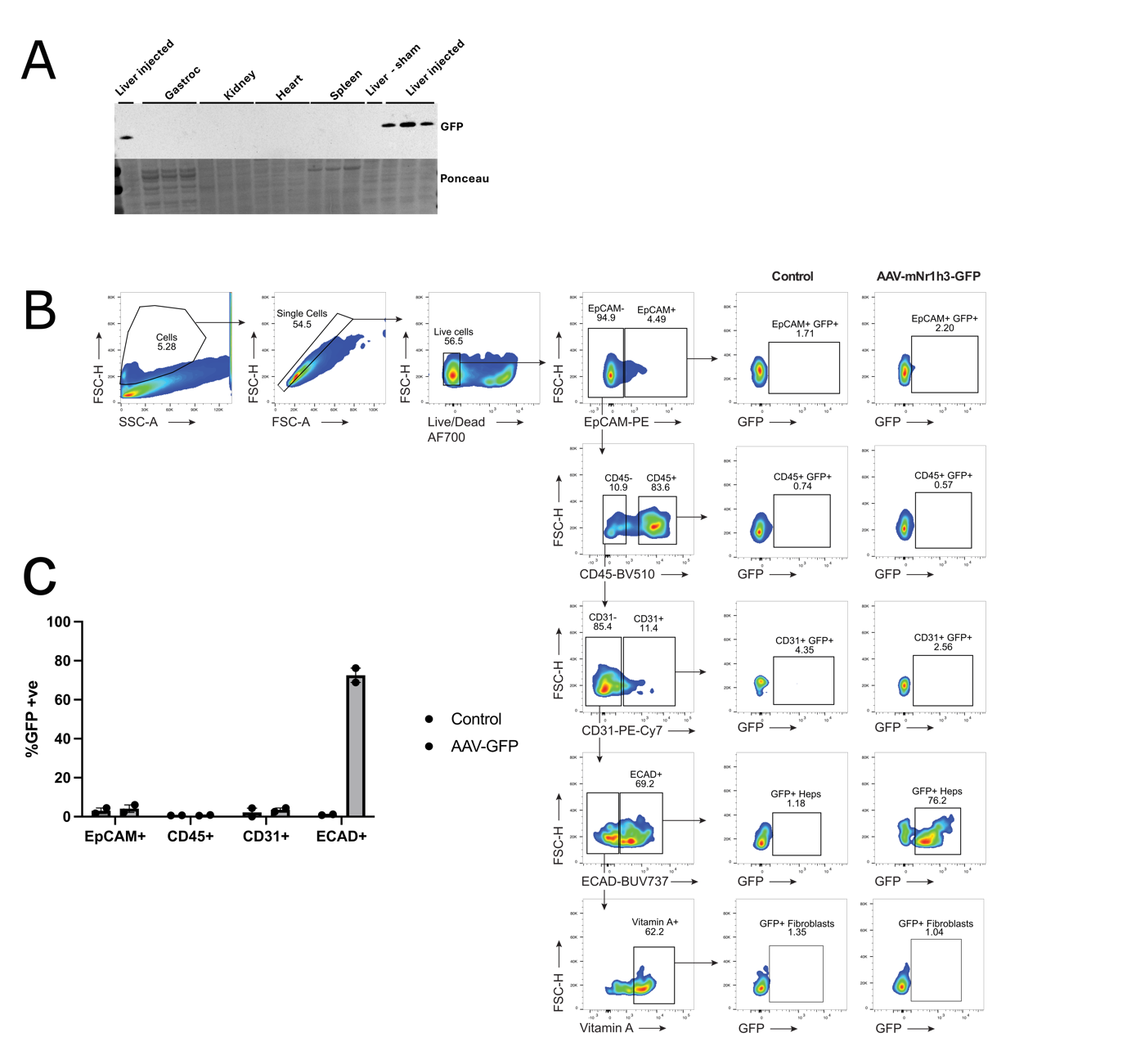

**Supplementary Figure 11. Validation of viral overexpression paradigm.** To validate that AAV8-mediated overexpression of wildtype *Nr1h3* was specific we detected the GFP marker gene transcribed from the bicistronic vector which also expressed *Nr1h3* using Western blot (A) and Flow Cytometry (B). **A:** GFP expression in a panel of selected tissue lysates was analysed by Western blot (N=3). A sample from a mouse that was subject to a sham injection of AAV8 is shown as a negative control, all other samples are from separate mice injected with AAV8-TBG-*Nr1h3*. **B:** Gating strategy for flow cytometry analysis of a single cell suspension from livers of mice subject to a sham injection or injected with AAV8-TBG-*Nr1h3*  (N=2 mice per group) which expresses marker eGFP gene from the same bicistronic vector as *Nr1h3* (AAV-mNr1h3-GFP). GFP-positive cells are indicative of cellular transduction and is restricted to hepatocytes. **C:** The proportion of GFP-positive cells in each lineage are shown (N=2).

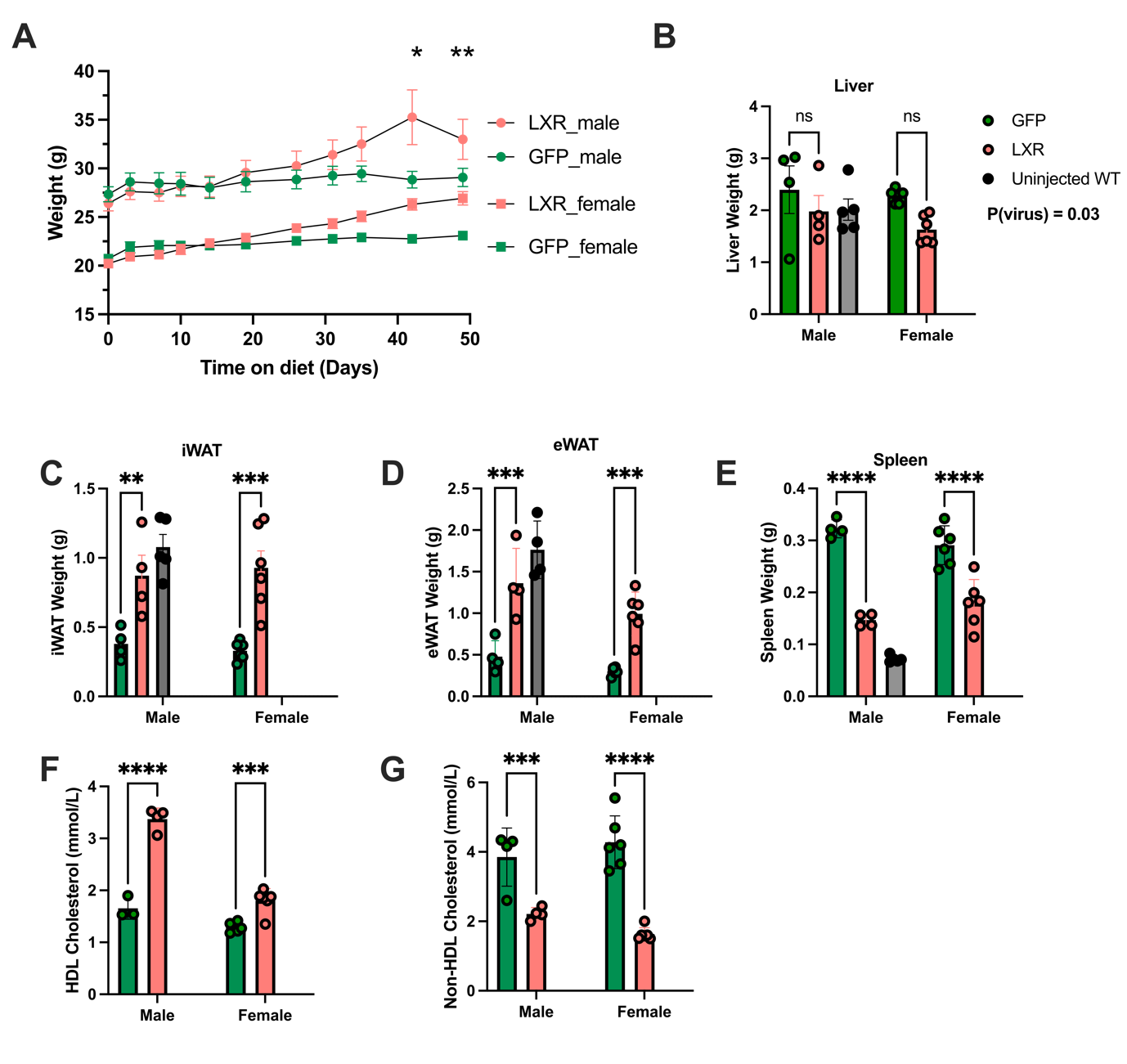

**Supplementary Figure 12 - Hepatocyte expression of WT mouse LXRα normalises bodyweight, organ weights and serum biochemistry in Western diet-fed LXRα^W441R/W441R^**  **mice.** 10 male and 12 female 10-12 week old C57BL/6J LXRα ^W441R /W441R^  mice were randomised to receive a tail vein injection with 1x10^-11^ GC of AAV8 expressing either LXRα and GFP (LXR) or just GFP (GFP) under the control of the hepatocyte specific Thyroxine binding globulin (TBG) promoter (see methods) and a week later they were placed on western diet. **A:** Body weight curves of mice in each group on a western diet. **B-E:** Tissues weights of indicated organs at necropsy **F-G:** Serum cholesterol measurements after 8 weeks on Western diet. Throughout, the red bars and symbols represent mice treated with LXRα expressing virus, green represents mice injected with control GFP-expressing virus, grey indicate non-littermate 10-week old male wildtype mice not injected with virus as a comparison (N=5). Repeated measures data (bodyweight) was analysed with a mixed effects model with post-hoc between group comparisons undertaken with the Holm-Šídak test with sex included in the model as a co-variate. All other data presented are analysed with a Two-way ANOVA with post-hoc  Holm-Šídak testing. Height of the bars represents mean +/- SEM.
